# *Bartonella* effector protein C mediates actin stress fiber formation via recruitment of GEF-H1 to the plasma membrane

**DOI:** 10.1101/2020.04.17.046482

**Authors:** Simon Marlaire, Christoph Dehio

## Abstract

*Bartonellae* are Gram-negative facultative-intracellular pathogens that use a type-IV-secretion system (T4SS) to translocate a cocktail of *Bartonella* effector proteins (Beps) into host cells to modulate diverse cellular functions. BepC was initially reported to act in concert with BepF in triggering major actin cytoskeletal rearrangements that result in the internalization of a large bacterial aggregate by the so-called ‘invasome’. Later, infection studies with *bepC* deletion mutants and ectopic expression of BepC have implicated this effector in triggering an actin-dependent cell contractility phenotype characterized by fragmentation of migrating cells due to deficient rear detachment at the trailing edge, and BepE was shown to counterbalance this remarkable phenotype. However, the molecular mechanism of how BepC triggers cytoskeletal changes and the host factors involved remained elusive. Using infection assays, we show here that T4SS-mediated transfer of BepC is sufficient to trigger stress fiber formation in non-migrating epithelial cells and additionally cell fragmentation in migrating endothelial cells. Interactomic analysis revealed binding of BepC to a complex of the Rho guanine nucleotide exchange factor GEF-H1 and the serine/threonine-protein kinase MRCKα. Knock-out cell lines revealed that only GEF-H1 is required for mediating BepC-triggered stress fiber formation and inhibitor studies implicated activation of the RhoA/ROCK pathway downstream of GEF-H1. Ectopic co-expression of tagged versions of GEF-H1 and BepC truncations revealed that the C-terminal ‘<u>B</u>ep intracellular <u>d</u>elivery’ (BID) domain facilitated anchorage of BepC to the plasma membrane, whereas the N-terminal ‘filamentation induced by <u>c</u>AMP’ (FIC) domain facilitated binding of GEF-H1. While FIC domains typically mediate post-translational modifications, most prominently AMPylation, a mutant with quadruple amino acid exchanges in the putative active site indicated that the BepC FIC domain acts in a non-catalytic manner to activate GEF-H1. Our data support a model in which BepC activates the RhoA/ROCK pathway by re-localization of GEF-H1 from microtubules to the plasma membrane.

**Author Summary:** A wide variety of bacterial pathogens evolved numerous virulence factors to subvert cellular processes in support of a successful infection process. Likewise, bacteria of the genus *Bartonella* translocate a cocktail of effector proteins (Beps) via a type-IV-secretion system into infected cells in order to interfere with host signaling processes involved in cytoskeletal dynamics, apoptosis control, and innate immune responses. In this study, we demonstrate that BepC triggers actin stress fiber formation and a linked cell fragmentation phenotype resulting from distortion of rear-end retraction during cell migration. The ability of BepC to induce actin stress fiber formation is directly associated with its ability to bind GEF-H1, an activator of the RhoA pathway that is sequestered in an inactive state when bound to microtubules, but becomes activated upon release to the cytoplasm. Our findings suggest that BepC is anchored via its BID domain to the plasma membrane where it recruits GEF-H1 via its FIC domain, eventually activating the RhoA/ROCK signaling pathway and leading to stress fiber formation.

## Introduction

The cytoskeleton plays major roles in epithelial and endothelial barrier integrity, pathogen uptake, and immune cell functions such as phagocytosis and cell migration. Depending on their infection strategies, pathogenic bacteria have evolved a plethora of virulence factors to obstruct or subvert these cytoskeletal functions. Many of these virulence factors target Rho family GTPases due to their central roles in regulating cytoskeletal dynamics. These virulence factors stimulate, attenuate or inactivate the intrinsic GTPase activities of Rho family GTPases, either directly through covalent modification [1], or indirectly by deregulating the activities of guanine nucleotide exchange factors (GEFs) [2] or GTPase-activating proteins (GAPs), or by molecular mimicry of GEF or GAP functions [3]. These virulence factors can be toxins, which are secreted to the extracellular milieu and are enabled to autonomously enter cells in order to reach their targets, or they are effector proteins, which are directly translocated into host cell via dedicated delivery devices, such as the type III (T3SS) or type IV (T4SS) secretion systems [4].

The gram-negative, facultative intracellular pathogens of the genus *Bartonella* are arthropod-borne bacteria that cause long-lasting intraerythrocytic bacteremia as hallmark of chronic infection in their specific mammalian reservoirs. While only few species are human-specific (e.g., *B. quintana*), many of the animal-specific species are zoonotic as they cause incidental human infections, resulting in a broad spectrum of clinical manifestations that ranges from asymptomatic courses to life-threatening disease [5]. For instance, the zoonotic pathogen *B. henselae* (*Bhe*) naturally infects cats, but is responsible for the majority of humans cases of *Bartonella* infection due to transmission to humans by cat scratch or bite. Infected immunocompetent individuals develop so-called cat scratch disease that leads to lymphadenopathy and fever, while immunocompromised patients develop bacillary angiomatosis characterized by vasoproliferative tumors of the skin and inner organs [6].

The *bartonellae* utilize a VirB/VirD4 T4SS to translocate a cocktail of *<u>B</u>artonella* <u>e</u>ffector <u>p</u>roteins (Beps) into host cells, and their orchestrated activities modulate multiple cellular processes and thereby decisively contribute to the stealth infection strategy and capacity of these pathogens to cause chronic infection [6, 7]. Beps are multi-domain proteins that share a common architecture at their C-terminus, which is composed of a ‘<u>B</u>ep intracellular <u>d</u>elivery’ (BID) domain and a positively charged tail that together constitute an evolutionary conserved bipartite signal for T4SS-mediated translocation [8, 9]. Despite their conserved fold [10], BID domains display significant variability in surface-exposed amino acids that facilitated the evolution of specific, non-enzymatic effector functions within host cells, e.g. by mediating protein-protein interaction or anchorage to the plasma membrane [11-15]. The N-terminus is more divergent. It may encode additional BID domains [13, 16], or tandem-repeated tyrosine phosphorylation motifs that serve as scaffolds for the assembly of signaling complexes [5, 13, 16-18], however, most Beps carry an N-terminal ‘filamentation induced by <u>c</u>AMP’’ (FIC) domain and a central OB (oligosaccharide binding) fold [16]. This conserved FIC-OB-BID domain order is also considered to represent the architecture of the ancestral effector from which all present-day Beps have evolved by gene duplication, domain shuffling, and sequence diversification [16, 17, 19]. FIC domains are characterized by a core composed of six α-helices, which includes a signature sequence called ‘FIC motif’ and a so-called ‘flap’ sequence [20, 21]. The canonical FIC motif HxFx(D/E)GNGRxxR plays a key role in the transfer of a diphosphate-containing group onto the hydroxyl side chain of the amino acids threonine (T), serine (S), or tyrosine (Y) in target proteins. Most FIC domains mediate the transfer of an AMP moiety from ATP as substrate by a reaction known as AMPylation or adenylylation, however, some FIC proteins are able to utilize different substrates to catalyze other posttranslational modifications [20, 21]. The flap of the FIC domain overlays the active site and mediates β-strand augmentation with the amino acid chain of target proteins to register a hydroxyl side-chain for modification [21, 22]. The OB fold connects the N-terminal FIC domain and the C-terminal BID domain. It may primarily serve as an interdomain fold [23], but despite its small size it may extend a protein-protein interaction interface and/or effector localization sequence composed by the proximal FIC or BID domains.

The *Bartonella* effector protein C (BepC) was reported to trigger two distinct F-actin driven cytoskeletal processes that are both dependent on actin stress fiber formation and dynamics [7]. First, BepC_*Bhe*_ was shown to act in concert with BepF_*Bhe*_ in triggering pronounced actin cytoskeletal rearrangement in primary <u>h</u>uman <u>u</u>mbilical <u>v</u>ein <u>e</u>ndothelial <u>c</u>ells (HUVECs) and epithelial HeLa cells that resulted in the internalization of a large bacterial aggregate by a multi-step process known as ‘invasome-mediated internalization’ [12, 24-26]. Then, infection assays with Δ*bepC*_*Bhe*_ deletion mutants in dendritic cells and HUVECs and ectopic expression of mCherry-BepC_*Bhe*_ in HUVECs implicated BepC in triggering actin stress fiber formation, resulting in the fragmentation of migratory cells due to deficient rear detachment at the trailing edge [11]. This remarkable phenotype based on imbalanced formation and disassembly of focal adhesion complexes during actomyosin-dependent cell contraction is at least in part antagonized by the activity of BepE [11]. On the structural level, BepC displays the ancestral FIC-OB-BID architecture. However, unlike most Fic proteins, BepC is characterized by a non-canonical but well conserved FIC motif (HxFx**K**GNGRxxR), which differs from the canonical motif by the replacement of an acidic residue (D/E) by a lysine (K). The crystal structure of the FIC domain of BepC from *Bartonella tribocorum* (BepC_*Btr*_), co-crystallized with an ATP derivative in the active site, indicated that the lysine is directly interacting with the α- and β-phosphates of the ATP analog (PDB: 4WGJ), thus functionally replacing the magnesium cation (Mg^2+^) that is coordinated by the acidic residue (D/E) of the canonical FIC motif [27]. Although this arrangement might be compatible with an AMP-transferase activity as shown for FIC domains with a conserved canonical FIC motif, no enzymatic activity has been reported yet for the BepC FIC domain. The BID domain of BepC_*Bhe*_ was shown to mediate effector translocation via the VirB/VirD4 T4SS [8] and to associate with the plasma membrane within host cells [14]. Despite these insights from structure/function analysis, the molecular mechanism of how BepC triggers cytoskeletal changes that contribute to invasome formation and cell fragmentation of migratory cells remained elusive, and no host targets of BepC have been reported to date.

In this study, we demonstrate that BepC triggers actin stress fiber formation by activating the RhoA GTPase signaling cascade via recruitment of GEF-H1 to the plasma membrane. We further show that BepC binds GEF-H1 via the FIC domain while anchorage to the plasma membrane depends on the BID domain.

## Results

### BepC triggers actin stress fiber formation and cell fragmentation in human umbilical vein endothelial cells

Infection with *Bhe* Δ*bepC*_*Bhe*_ deletion mutants and ectopic expression of mCherry-BepC_*Bhe*_ in HUVECs implicated BepC in triggering actin stress fiber formation and the linked cell fragmentation phenotype resulting from distorted rear-end detachment during cell migration [11]. To demonstrate that these prominent phenotypes can result from VirB/VirD4-dependent translocation of BepC_*Bhe*_ into infected HUVECs we expressed FLAG-BepC_*Bhe*_ (Fig. 1A) in the effector-free background of the *Bhe* Δ*bepA-G* strain [8]. Translocation of FLAG-BepC_*Bhe*_ by this strain triggered both F-actin-dependent phenotypes in dependency of infection time and multiplicity of infection (MOI) (Fig. 1B and Fig. S1), while the isogenic control strain containing the empty expression plasmid did not noticeably alter the F-actin cytoskeleton compared to uninfected cells (Fig. 1B and Fig. S1). BepC_*Bhe*_-dependent actin stress fiber formation was evident already at MOI of 50 at 24 h, while at this time-point cell fragmentation became only visible at MOI of 400. Generally, stress fiber formation and cell fragmentation phenotypes were more pronounced at 48 h than at 24 h, and this late time-point was thus used for most follow-up experiments. Of note, BepC-triggered cell fragmentation resulted in decreased cell number, although fragmented cells did not seem to undergo apoptosis or necrotic cell death.

**Fig 1.**
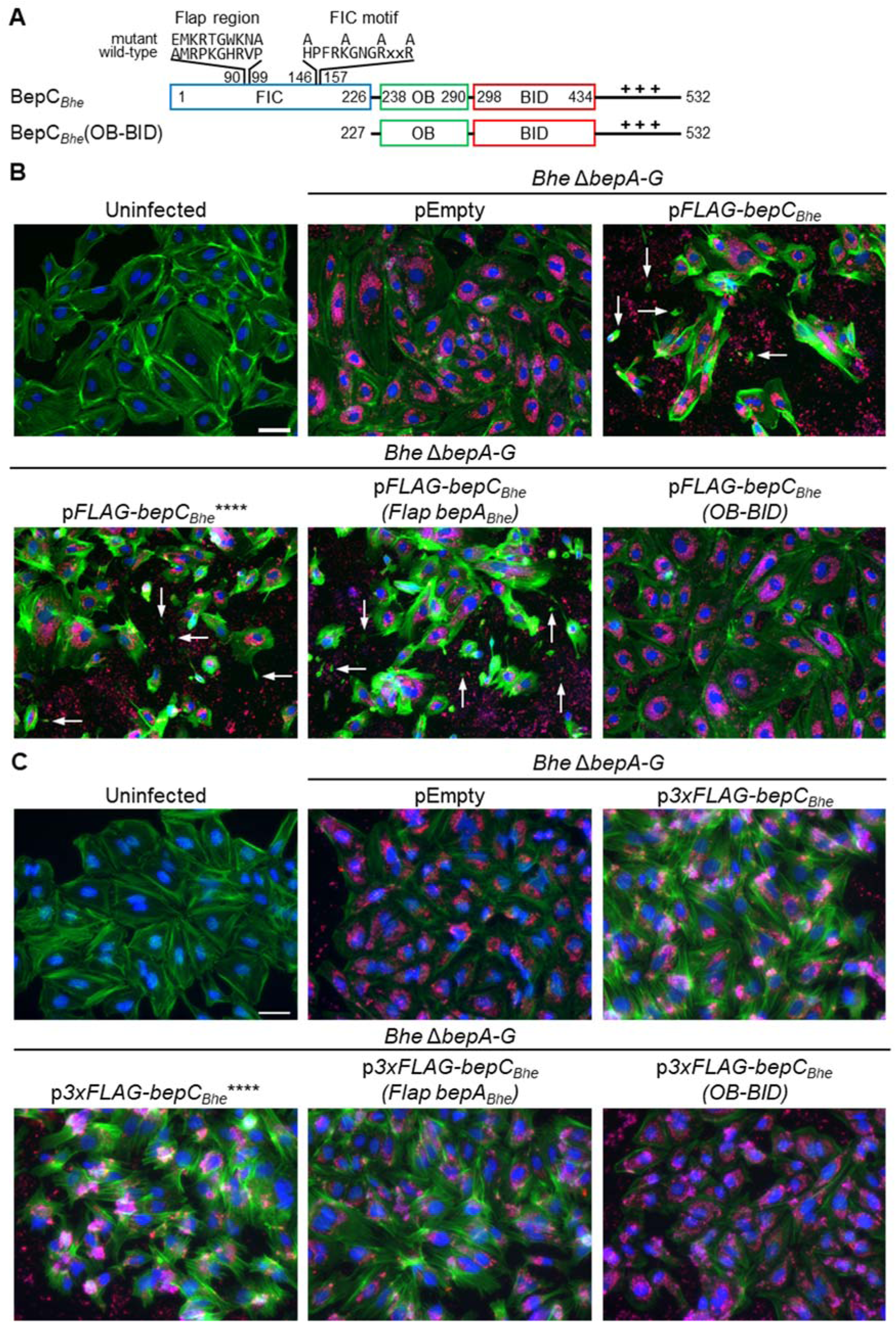
The BepC_*Bhe*_ FIC domain but not a conserved FIC motif or flap region is required for actin stress fiber formation in *B. henselae*-infected HUVECs or HeLa cells. (**A**) Schematic view of BepC_*Bhe*_ wild-type and mutant variants analyzed in this figure. The positively charged tail at the C-terminus is represented by +++. The N-terminally fused FLAG-tag in either singular or triple copy is not shown. (**B**-**C**) HUVECs (**B**) or HeLa cells (**C**) were infected with isogenic *Bhe* Δ*bepA-G* strains expressing FLAG-tagged BepC_*Bhe*_ wild-type or mutant versions, or carrying the empty plasmid at multiplicity of infection (MOI) of 400. After 48 h of infection, cells were fixed and immunocytochemically stained, followed by fluorescence microscopy analysis. F-actin is represented in green, DNA in blue, and bacteria in red (scale bar = 50 µm). Arrows point to cell fragments resulting from distorted rear end retraction of migrating HUVEC. BepC_*Bhe*_**** = BepC_*Bhe*_ H146A, K150A, R154A, R157A.

The slow kinetics and high MOI-dependency of BepC_*Bhe*_–triggered stress fiber formation suggest that the effector may act on its host target(s) by protein-protein interaction (e.g., by target sequestration or mislocalization) rather than by an enzymatic mechanism. While BepC contains a well-conserved FIC domain that typically catalyzes an enzymatic activity such as AMPylation [20], its FIC motif (HxFx**K**GNGRxxR) differs from the canonical FIC motif (HxFx**D/E**GNGRxxR), which is defined by essential amino acids in the active site of these AMP-transferases, in one amino acid position (D/E replaced by K) [16]. Despite this single amino acid exchange, the FIC domain of BepC might still harbor enzymatic activity; possibly one that differs from AMPylation. However, we reasoned that the mutations of this lysine and of three additional amino acids known to be essential for enzymatic activity in FIC domains with a canonical FIC motif [20, 21, 28] should incapacitate any presumable enzymatic activity of the BepC FIC domain. We thus constructed the quadruple mutant BepC_*Bhe*_**** with the amino acid exchanges H146A, K150A, R154A, R157A, resulting in a highly degenerated FIC motif (**A**xFx**A**GNG**A**xx**A**, Fig. 1A). Moreover, we constructed another mutant that might compromise BepC-specific target modification by exchanging the flap region, which registers the target protein to the active site of FIC domains, between BepC_*Bhe*_ and BepA_*Bhe*_ (FLAG-BepC_*Bhe*_(Flap bepA_*Bhe*_), Fig. 1A). Both of these BepC_*Bhe*_ mutant proteins maintained the same capacity to trigger stress fiber formation as BepC_*Bhe*_, indicating that BepC_*Bhe*_ may not require enzymatic modification of host targets to trigger actin rearrangements (Fig. 1B). However, deletion of the FIC domain (FLAG-BepC_*Bhe*_(OB-BID), Fig. 1A) rendered the truncated BepC_*Bhe*_ protein inactive (Fig. 1B).

In summary, the FIC domain of BepC is required for actin stress fiber formation, but neither the conserved FIC motif nor the specific flap region are necessary, suggesting that the actin phenotype is linked to a non-enzymatic activity.

### BepC-triggered actin stress fiber formation in HeLa cells is dependent on both the FIC and the BID domain

To facilitate further cellular and molecular analysis of the mechanism underlying BepC-triggered actin stress fiber formation in an established cell line we adopted the previously published HeLa cells infection model [25]. To facilitate improved FLAG-tag-based pull-down of BepC variants in subsequent interactomic studies we generated triple FLAG-tag fusions for HeLa infection (Fig. 1C) instead of the singular FLAG-tag fusions used for HUVEC infection (Fig. 1B). This minor change in the protein tagging strategy had no functional consequences given that the capacity of isogenic strains expressing BepC_*Bhe*_ wild-type and corresponding mutant variants to trigger actin stress fiber formation in HUVEC was fully reproduced in HeLa cells (Fig. 1C).

As a complementary approach to VirB/VirD4-dependent effector translocation, we tested how ectopic expression of BepC_*Bhe*_ wild-type and mutant variants affected actin stress fiber formation. The phenotypic data obtained for ectopic expression in HeLa cells (Fig. 2) are in full agreement with translocation-dependent infection phenotypes in both HeLa and HUVEC (Fig. 1B, C). Importantly, the ectopic expression approach allowed us to test also a C-terminal truncation resulting in deletion of the entire BID domain and positively charged tail sequence (3xFLAG-BepC_*Bhe*_(FIC-OB), Fig. 2A) that could not be tested in the infection assay as deletion of this bipartite secretion signal abolishes VirB/VirD4-dependent protein translocation [8]. The lack of increased actin stress fiber formation by ectopic expression of this C-terminal truncation demonstrated the essential role of the BID domain in mediating this phenotype (Fig. 2B).

**Fig 2.**
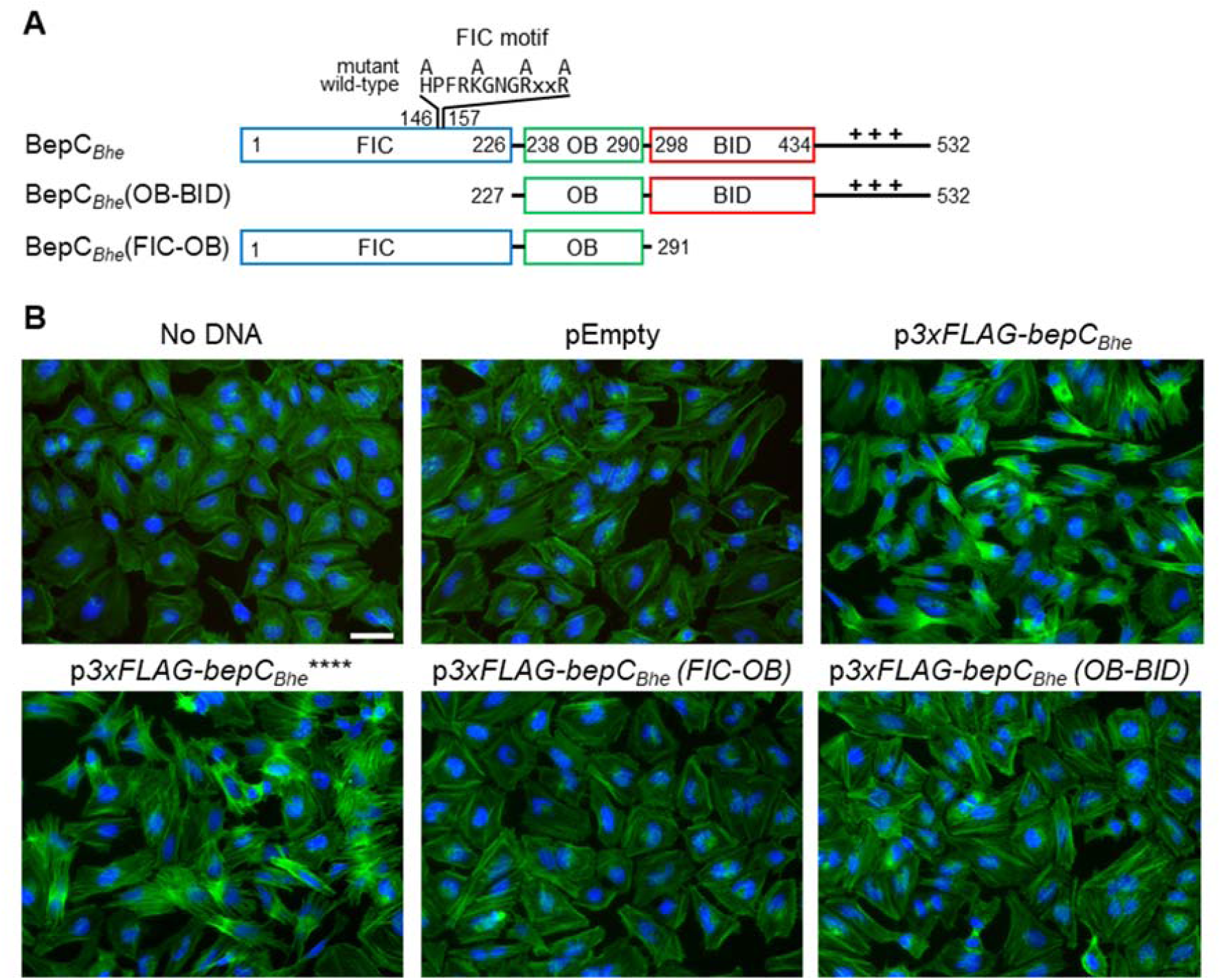
Both FIC and BID domains are required for BepC_*Bhe*_–triggered actin stress fiber formation upon ectopic effector expression in HeLa cells. (**A**) Schematic view of BepC_*Bhe*_ wild-type and mutant variants analyzed in this figure. The positively charged tail at the C-terminus is represented by +++. The N-terminally fused triple FLAG-tag is not shown. (**B**) HeLa cells were transfected with the indicated plasmids for expression of FLAG-tagged BepC_*Bhe*_ wild-type, mutant versions, or truncations, or no protein as negative control (pEmpty). 24 h after transfection, cells were fixed and immunocytochemically stained, followed by fluorescence microscopic analysis. F-actin is represented in green and DNA in blue (scale bar = 50 µm). BepC_*Bhe*_**** = BepC_*Bhe*_ H146A, K150A, R154A, R157A.

The high level of sequence conservation of BepC homologs in different *Bartonella* species is indicative of a conserved molecular function [16]. We thus tested whether the capacity of BepC_*Bhe*_ to trigger actin stress fiber formation is conserved among BepC homologs. HeLa cells were infected with *Bhe* Δ*bepA-G* expressing FLAG-tagged BepC of *Bhe, B. quintana, B. tribocorum, B. taylorii*, or *B. grahamii*. Increased F-actin stress fiber formation was evident for all BepC homologs, except for BepC_*Bgr*_ that was indistinguishable from the negative control strain *Bhe* Δ*bepA-G* pEmpty (Fig. S2A). However, ectopic expression of these natural BepC variants demonstrated strongly increased stress fiber formation for each of them, including BepC_*Bgr*_ (Fig. S2B), suggesting that the lack of phenotype for BepC_*Bgr*_ in the infection assay may be a false-negative result, possibly because of insufficient FLAG-BepC_*Bgr*_ expression, stability, or translocation.

In summary, our data demonstrate that both FIC and BID domains are required for BepC-triggered actin stress fiber formation. Moreover, these actin rearrangements are triggered by all tested BepC homologs from various *Bartonella* species, suggesting that this conserved effector function targeting the actin cytoskeleton is likely playing a crucial role during *Bartonella* infection.

### GEF-H1 and MRCKα form a complex that co-immunoprecipitates with BepC

In order to search for potential host targets of BepC_*Bhe*_, we identified interacting host proteins by an interactomics approach. To this end, we infected HeLa cells with *Bhe* Δ*bepA-G* expressing FLAG-tagged BepC_*Bhe*_ or the isogenic strain containing the empty expression plasmid. Following cell lysis, 3xFLAG-BepC_*Bhe*_ was pulled down with anti-FLAG-tag antibodies and co-immunoprecipitating host proteins were identified by mass spectrometry (Fig. 3A and Table S1). Compared to the negative control, six proteins showed an increase of at least 8-fold and a q-value lower than 0.05 (Fig. 3A). Three of these six proteins were identified by a single peptide. These proteins were not followed up as their relevance was uncertain. As expected, one of the three remaining proteins corresponded to BepC_*Bhe*_ as the bait. The two outstanding interactors, GEF-H1 and MRCKα, were particularly interesting as both are involved in regulating F-actin rearrangements (Fig. 4A). On one hand, GEF-H1 promotes the activation of RhoA by exchanging GDP for GTP, which then via ROCK activation and subsequent phosphorylation of myosin light chain (MLC) ultimately leads to actin stress fiber formation [28-31] (Fig. 4A). On the other hand, MRCKα is a downstream effector of the Cdc42 pathway and directly phosphorylates MLC and, as ROCK, inhibits the myosin light chain phosphatase (MLCP), thereby promoting actin stress fiber formation as well [29-31] (Fig. 4A).

**Fig 3.**
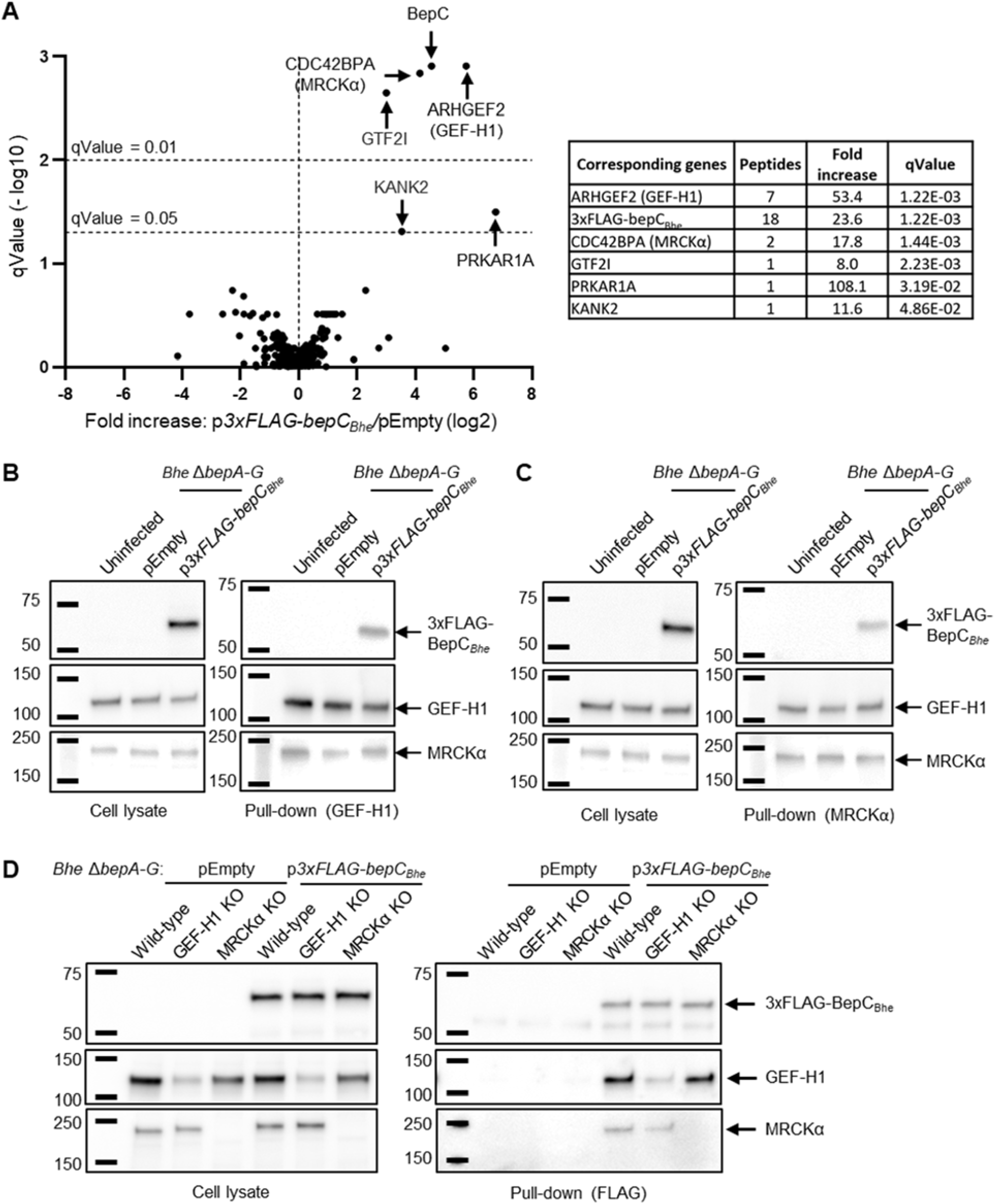
BepC_*Bhe*_ binds directly to GEF-H1 in a trimeric complex with MRCKα. (**A**) HeLa cells were infected with *Bhe* Δ*bepA-G* expressing FLAG-tagged BepC_*Bhe*_, or carrying the empty plasmid as a negative control, at MOI of 200. After 24 h of infection, cells were lysed and the lysate incubated in presence of anti-FLAG antibody. 3xFLAG-BepC_*Bhe*_ and interacting proteins were pulled-down with protein G agarose beads and bound proteins were released with SDS-containing buffer. Samples (technical triplicates) were analyzed by mass spectrometry and data obtained for 3xFLAG-BepC_*Bhe*_ and the negative control were compared (see Table S1 for a listing of all identified proteins). (**B**-**C**) HUVECs were infected with *Bhe* Δ*bepA-G* expressing 3xFLAG-tagged BepC_*Bhe*_ or carrying empty plasmid at MOI of 200 for 24 h. Cells were lysed and incubated in presence of (**B**) anti-GEF-H1 antibody or (**C**) anti-MRCKα antibody. Antibody-bound proteins were subsequently pulled-down with protein G agarose beads, followed by elution with SDS-containing buffer. Cell lysates before pull-down and pull-down samples were analyzed by immunoblot using antibodies against FLAG-tag, GEF-H1, or MRCKα. (**D**) HeLa wild-type or knocked-out cells for GEF-H1 or MRCKα were infected with *Bhe* Δ*bepA-G* expressing FLAG-tagged BepC_*Bhe*_ or carrying the empty plasmid as a negative control at MOI of 200. After 24 h of infection, cells were lysed and incubated with anti-FLAG antibodies. 3xFLAG-BepC_*Bhe*_ was pulled-down with protein G agarose beads before eluted with SDS. Cell lysates before pull-down and pull-down samples were analyzed by immunoblot against FLAG-tag, GEF-H1, or MRCKα.

**Fig 4.**
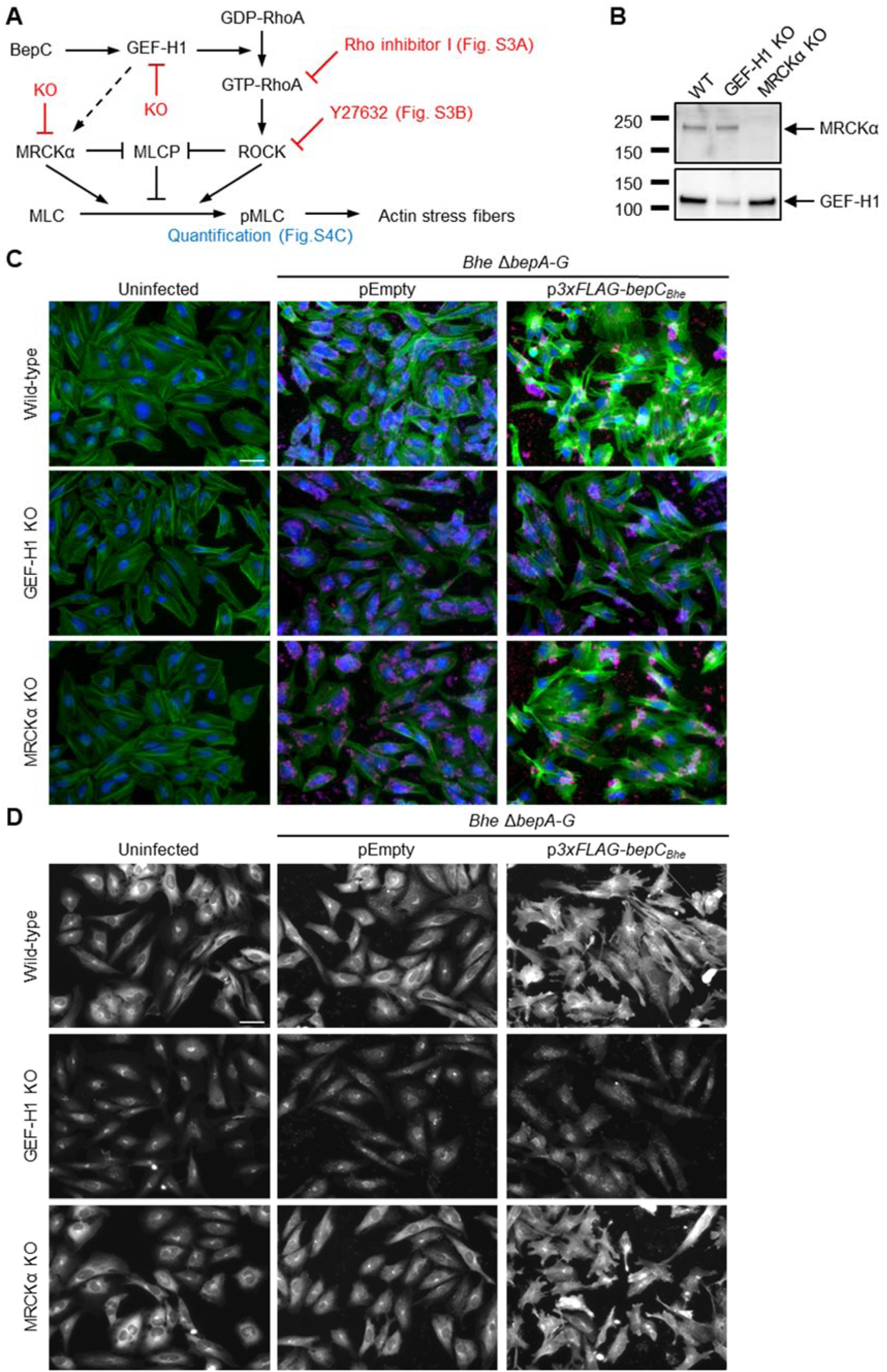
GEF-H1 is essential for BepC_*Bhe*_–triggered actin stress fiber formation while MRCKα is dispensable. (**A**) Proposed model of BepC-triggered actin stress fiber formation with reference to the experimental data presented for validation. (**B**) HeLa cells were co-transfected with two different plasmids encoding Cas9 and a sgRNA, specific either to the first or the last exon of the target gene (GEF-H1 or MRCKα). After selection and expansion of transfected cells, expression of GEF-H1 or MRCKα was tested by immunoblot analysis. (**C-D**) HeLa cells wild-type, GEF-K1 KO, and MRCKα KO were infected with *Bhe* Δ*bepA-G* expressing FLAG-tagged BepC_*Bhe*_, or carrying the empty plasmid as a negative control, at MOI of 400. After 48 h of infection, cells were fixed, stained by immunocytochemistry and analyzed by fluorescence microscopy. (**C**) F-actin is represented in green, DNA in blue, and bacteria in red. (**D**) GEF-H1 is represented in white (scale bar = 50 µm).

To validate the BepC_*Bhe*_ interaction with GEF-H1 and MRCKα as identified by interactomics in HeLa cells we performed a reciprocal co-immunoprecipitation experiment in a different cell type. To this end, HUVECs were infected with *Bhe* Δ*bepA-G* expressing FLAG-tagged BepC_*Bhe*_ or carrying the empty expression plasmid. Following pull-down of GEF-H1 (Fig. 3B) or MRCKα (Fig. 3C), 3xFLAG-BepC_*Bhe*_ was detected in both pull-down fractions, confirming that the three proteins are forming a trimeric complex. However, even in the absence of BepC_*Bhe*_ GEF-H1 co-immunoprecipitated with MRCKα and vice versa, indicating that MRCKα and GEF-H1 are part of a native complex that to our knowledge has not been described yet (Fig. 3B,C).

In summary, interactomics identified a native complex of GEF-H1 and MRCKα that co-immunoprecipitated with BepC_*Bhe*_, leaving it open whether BepC interacts in a direct manner with GEF-H1, or with MRCKα, or with both.

### BepC binds directly to GEF-H1 in a trimeric complex with MRCKα

To distinguish between direct and indirect interactions between BepC_*Bhe*_ and either GEF-H1 or MRCKα, we generated HeLa knock-out cell lines for these cellular targets by CRISPR/Cas9 gene editing. To this end, we co-transfected HeLa cells with two different plasmids encoding Cas9 and sgRNAs that are specific to either the first or the last exon of the gene of interest. Following selection and polyclonal expansion of knock-out cells, we tested expression of GEF-H1 or MRCKα via immunoblot analysis. The data indicate a complete knock-out for MRCKα, and a partial knock-out for GEF-H1 with a nevertheless strongly diminished protein level compared to the parental wild-type cell line (Fig. 4B).

The established knock-out cell lines for GEF-H1 or MRCKα and the parental wild-type cell line were then used to untangle the molecular interactions within the trimeric complex of BepC, GEF-H1, and MRCKα via infection and co-immunoprecipitation analysis. To this end, the three cell lines were infected with *Bhe* Δ*bepA-G* expressing FLAG-tagged BepC_*Bhe*_ or carrying the empty expression plasmid. Following cell lysis, 3xFLAG-BepC_*Bhe*_ was pulled-down and tested for co-immunoprecipitation of GEF-H1 and/or MRCKα by immunoblot analysis (Fig. 3D). GEF-H1 interaction with 3xFLAG-BepC_*Bhe*_ was indistinguishable for wild-type cells and MRCKα knock-out cells, indicating that the interaction must be direct and independent of MRCKα (Fig. 3D, pull-down). In contrast, MRCKα interaction with 3xFLAG-BepC_*Bhe*_ was reduced in the partial GEF-H1 knock-out cell line compared to wild-type cells, suggesting that this interaction might be indirect. Clarification of this finding may require the establishment of a complete GEF-H1 knock-out cell line, which, however, may not be viable.

Overall, our data indicate that BepC_*Bhe*_ interacts directly with GEF-H1, while we could not unambiguously resolve the exact nature of the interaction with MRCKα, which thus may occur either directly, or indirectly through binding to GEF-H1.

### BepC–triggered actin stress fiber formation requires GEF-H1 and is associated with GEF-H1 relocalization

Next, we used the established GEF-H1 or MRCKα knock-out cells and parental wild-type cells in infection experiments to test for the roles of GEF-H1 and MRCKα in mediating BepC_*Bhe*_–triggered stress fiber formation (Fig. 4A). To this end, we infected these three cell lines with *Bhe* Δ*bepA-G* expressing FLAG-tagged BepC_*Bhe*_ or carrying the empty expression plasmid, or left them uninfected as an additional negative control. Staining for F-actin unequivocally demonstrated that GEF-H1 is essential for mediating BepC_*Bhe*_–triggered stress fiber formation, while MRCKα is neglectable for this process (Fig. 4C). Strikingly, staining with anti-GEF-H1 antibodies provided first evidence for a BepC–dependent relocalization of GEF-H1 (Fig. 4D). The GEF-H1 knock-out cells displayed invariant background staining for all three conditions. However, parental wild-type and MRCKα knock-out cells displayed a characteristic cytoplasmic GEF-H1 staining pattern consistent with microtubular association in both uninfected and control infection conditions. In contrast, GEF-H1 seemed to relocalize to the plasma membrane upon infection with the FLAG-tagged BepC_*Bhe*_ expressing strain (Fig. 4D).

In summary, we demonstrated an essential role of GEF-H1 for BepC_*Bhe*_– dependent actin stress fiber formation and provided first indications that BepC_*Bhe*_ translocation mediates a relocalization of GEF-H1 from a canonical microtubular-association to a putative plasma membrane localization.

### BepC interacts with GEF-H1 via its FIC domain while plasma membrane association is mediated by the BID domain

To corroborate our findings on a BepC-dependent relocalization of GEF-H1 from microtubules to the plasma membrane and to determine which BepC domain interactions are involved in mediating this effect, we have used an ectopic co-expression approach in HeLa cells. To follow GEF-H1 localization, we expressed a functional eGFP-GEF-H1 fusion that was previously reported to localize primarily to microtubules, indistinguishably from endogenous GEF-H1 [32]. Co-transfection with an empty expression plasmid (pEmpty) confirmed this canonical staining pattern as demonstrated by colocalization with microtubules (Fig. 5A). However, co-expression with Flag-tagged BepC_*Bhe*_ showed that part of the GEF-H1 signal remained associated with microtubules, while a significant proportion of the eGFP-GEF-H1 signal co-localized with 3xFLAG-BepC_*Bhe*_ at the plasma membrane (Fig. 5B, upper panel of x-y projections and left panel of x-z sections). In sharp contrast, the microtubule-associated localization of eGFP-GEF-H1 was unperturbed when co-expressed with either 3xFLAG-BepC_*Bhe*_(FIC-OB) or 3xFLAG-BepC_*Bhe*_(OB-BID) truncation constructs (Fig. 5B, middle or lower panel of x-y projections or middle or right panel of x-z sections, respectively). Strikingly, 3xFLAG-BepC_*Bhe*_(FIC-OB) perfectly colocalized with both eGFP-GEF-H1 and microtubules (Fig. 5B, middle panels of x-y projections and x-z sections), indicating that the soluble FIC-OB fragment binds to GEF-H1 without dissociating it from microtubules. On the contrary, 3xFLAG-BepC_*Bhe*_(OB-BID) displayed a plasma membrane localization without any sign of co-localization with microtubule-bound eGFP-GEF-H1 (Fig. 5B, lower panel of x-y projections and right panel of x-z sections).

**Fig 5.**
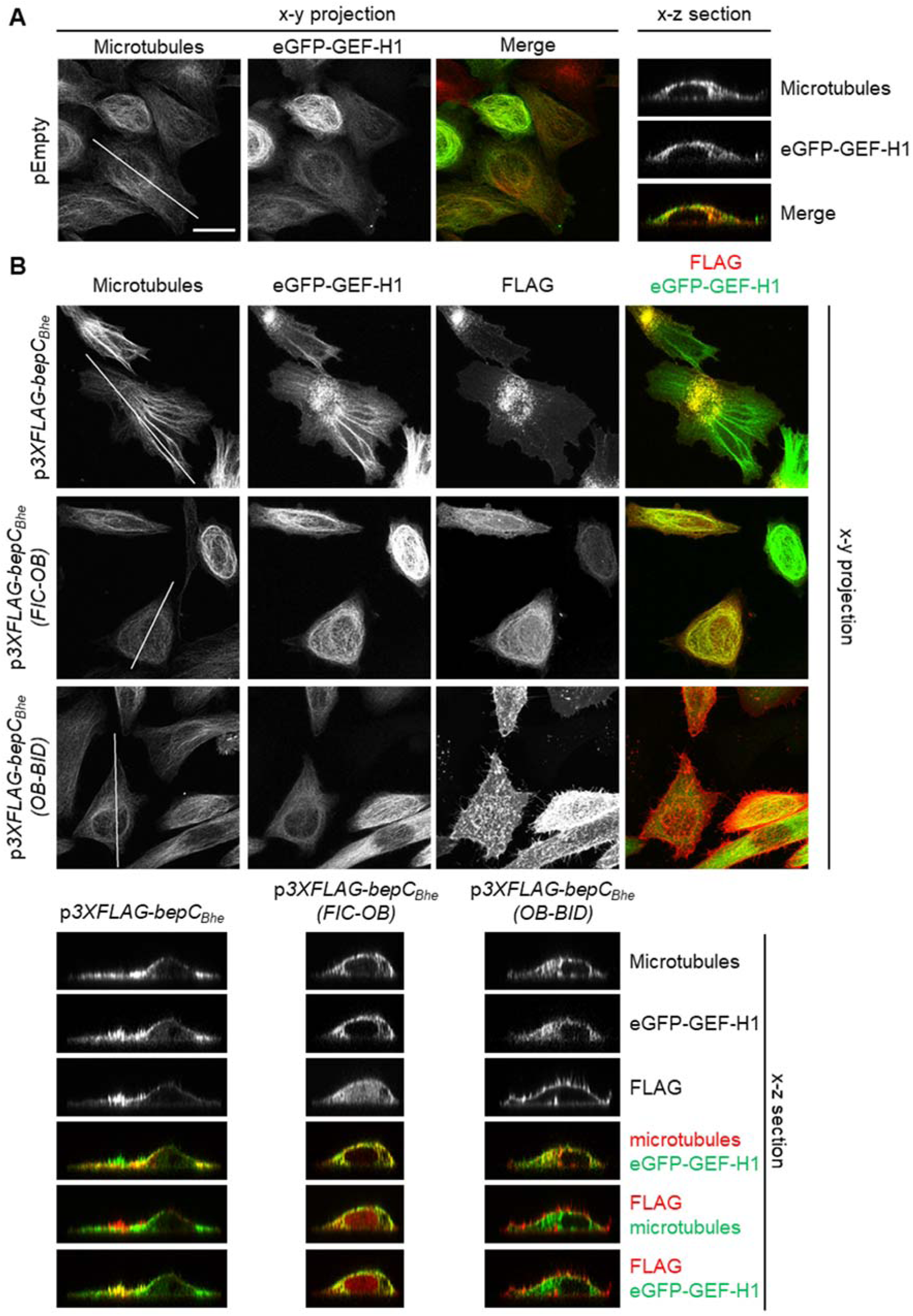
BepC_*Bhe*_ recruits eGFP-GEF-H1 to the plasma membrane via binding of the FIC-OB domain to eGFP-GEF-H1 and binding of the BID domain to the plasma membrane. (**A**-**B**) HeLa cells were co-transfected with an expression plasmid for eGFP-GEF-H1 and the indicated plasmids for either (**A**) no expression or (**B**) expression of either 3xFLAG-BepC_*Bhe*_, 3xFLAG-BepC_*Bhe*_(FIC-OB), or 3xFLAG-BepC_*Bhe*_(OB-BID). After 24 h, cells were fixed and stained by immunofluorescence labeling for FLAG and microtubule before being analyzed by fluorescence microscopy (scale bar = 25 µm). The x-z sections presented correspond to orthogonal cuts at the white lines displayed in the microtubule channel.

Taken together, these data strongly indicate that the FIC domain and possibly the OB fold is required for BepC binding to GEF-H1, while the BID domain and possibly the OB fold is necessary for plasma membrane interaction. Together, we conclude that these domain interactions mediate the BepC-dependent relocalization of GEF-H1 from the canonical microtubule-association to the plasma membrane.

### BepC-triggers actin stress fiber formation via the GEF-H1/RhoA/ROCK/pMLC pathway

While GEF-H1 binds to microtubules in an inactive conformation it gains GEF activity in association with membranes [32, 33], where it activates either RhoA or Rac1 [34]. BepC-mediated recruitment of GEF-H1 to the plasma membrane should thus activate the RhoA and/or Rac1 pathway. Given that the RhoA pathway triggers stress fiber formation, while the Rac1 leads to lamellipodia formation [35], we reasoned that BepC/GEF-H1-mediated stress fiber formation is dependent on the RhoA pathway. To demonstrate this experimentally, we tested the involvement of components of the RhoA pathway in the BepC-triggered phenotype, i.e., by inhibiting RhoA or the Rho-kinase ROCK, and by evaluating the phosphorylation of the ROCK-substrate myosin light chain (pMLC) (Fig. S3A and Fig. S4A). Rho inhibitor I, which inactivates RhoA, RhoB and RhoC in living cells [36], was found to interfere with stress fiber formation mediated by 3xFLAG-BepC_*Bhe*_ in a concentration dependent manner (Fig. S3B). Similarly, Y-27632, which inhibits ROCK in living cells [37], inhibited stress fiber formation by 3xFLAG-BepC_*Bhe*_ in a concentration-dependent manner (Fig. S3C). Finally, phosphorylation levels of pMLC correlated with stress fiber formation triggered by 3xFLAG-BepC_*Bhe*_ and active mutant constructs (Fig. S4).

Taken together, these data indicate that BepC activates a GEF-H1/RhoA/ROCK/pMLC signaling pathway in order to trigger actin stress fiber formation.

## Discussion

Manipulation of the host cell actin cytoskeleton is crucial for many bacterial pathogens in order to cross epithelial or endothelial barriers, to disseminate into deeper tissue sites, to invade non-phagocytic cells, or to prevent phagocytosis by professional phagocytes [38]. These pathogens have evolved numerous toxins and effector proteins that interfere with actin cytoskeletal dynamics, typically by modulating the activities of Rho GTPases. Frontal-attack pathogens that cause acute infection often encode potent virulence factors that target the entire cellular pool of Rho GTPases by covalent modification or via molecular mimicry of GEFs or GAPs [3]. In contrast, pathogens causing chronic infections may selectively target subsets of Rho GTPases, e.g. by modulating the activity of one of the many endogenous GEFs or GAPs that usually control the activity of only a small subset of Rho GTPases in a temporal and spatial manner, thereby mediating rather subtle cytoskeletal changes that are more compatible with their stealth-attack infection strategy [3]. Here, we demonstrate a new mechanism of targeted deregulation of an endogenous GEF that would be beneficial for stealth pathogens involved in chronic infections. We show that the T4SS-translocated effector BepC of *Bartonella* spp. recruits GEF-H1 to the plasma membrane, thereby activating the RhoA/ROCK signaling pathway and leading to actin rearrangements.

While some bacterial effectors induce microtubule depolymerization to release GEF-H1 and eventually activate the RhoA pathway [39], VopO, a T3SS effector from *Vibrio parahaemolyticus*, is the only bacterial effector reported to interact directly with GEF H1 and activate the RhoA pathway, thereby triggering actin stress fiber formation [2]. However, the mechanism of GEF H1 activation by VopO remained elusive as co-localization studies did not show an alteration of GEF-H1 localization or an increase of its GEF activity.

Given that BepC contains a FIC domain, which in the context of other Fic proteins is known to catalyze posttranslational modifications [20, 21, 28], it was conceivable to assume that the BepC-triggered actin phenotype might be associated with a putative enzymatic activity. However, the mutation of four essential amino acids in the conserved FIC motif of BepC_*Bhe*_ (H146A, K150A, R154A, R157A) or the exchange of the flap region required for registration of the target amino acid did not display any negative impact on actin stress fiber formation. Assuming that these modifications have compromised, if not fully eliminated the catalytic activity, we concluded that BepC acts likely on GEF-H1 via protein-protein interactions rather than by posttranslational modification. As the FIC motif is highly conserved between BepC homologs of different *Bartonella* species, it may still play another significant role during infection and we cannot exclude that BepC catalyzes a posttranslational modification on an unrelated target that is irrelevant for BepC-triggered actin stress fiber formation.

Interestingly, the unexpected finding that a FIC domain exerts a biological function unrelated to a catalytic activity is unique and only remotely reminiscent of the Fic protein AvrB, which lacks all residues required for enzymatic activity [40]. Thus, it opens new perspectives for the function of the many Beps, and other Fic proteins, carrying a non-canonical FIC motif and possibly also lacking enzymatic activity [16, 20, 41].

Ectopic expression of eGFP-tagged GEF-H1 together with FLAG-tagged BepC_*Bhe*_ full-length, or its FIC-OB or OB-BID truncation constructs, allowed us to develop a simple model of the activation of GEF-H1 by BepC (Fig. 6). BepC_*Bhe*_ full-length was found to localize to the plasma membrane and recruit GEF-H1 from its canonical microtubule-bound location. In sharp contrast, BepC_*Bhe*_(FIC-OB) co-localized with eGFP-GEF-H1 at microtubules, indicating that the soluble FIC domain binds to GEF-H1 without dissociating GEF-H1 from microtubules. On the contrary, OB-BID associated with the plasma membrane without any sign of co-localization with GEF-H1. Consistent with the latter finding, the BID domain of BepC_*Bhe*_ was previously reported to localize to the plasma membrane after ectopic expression in HEK293T cells [14]. Accordingly, ectopic expression of mCherry-tagged BepC_*Bhe*_ full-length in HUVECs was also reported to localize to the plasma membrane [11]. In conclusion, BepC_*Bhe*_ appears to bind GEF-H1 via the FIC domain and recruit it to the plasma membrane via anchorage by the BID domain (Fig. 6). Although further investigation is required to determine how GEF-H1 is recruited from the microtubule-bound state, we can formulate two hypotheses: i) Either membrane-bound BepC recruits over time the GEF-H1 sub-pool that is cycling between microtubules and the plasma membrane in the course of other signaling processes, or ii) only BepC full-length has the capacity to actively dissociate GEF-H1 from the microtubules and to relocalize it to the plasma membrane. In both cases, the GEF-H1 pool recruited to BepC at the plasma membrane should lead to activation of membrane-anchored RhoA via its GEF activity, followed by activation of the downstream Rho kinase ROCK, which in turn phosphorylates myosin light chain (MLC), eventually leading to actin stress fiber formation (Fig.6). Our data on Rho and ROCK inhibitors and phosphorylation of myosin light chain support this signaling cascade downstream of GEF-H1. In conclusion, we characterized a novel molecular mechanism by which bacterial pathogens may selectively activate a Rho GTPase pathway via the recruitment of GEF-H1 to the plasma membrane.

**Fig 6.**
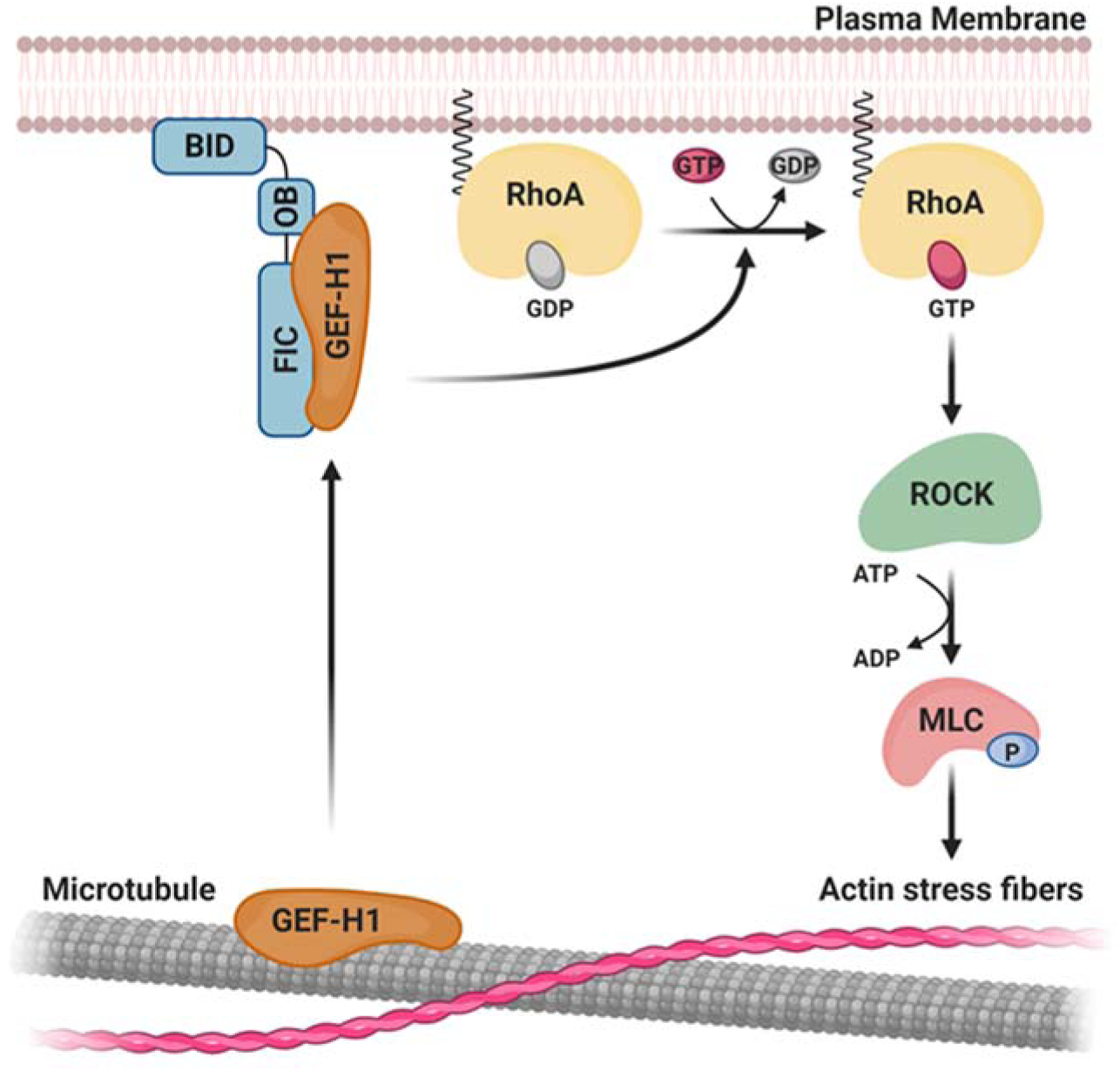
Model of BepC-triggered actin stress fibers formation mediated by the recruitment of GEF-H1 to the plasma membrane. Upon translocation, BepC localizes to the plasma membrane via its BID domain and binds GEF-H1 via its FIC-OB domains. There, GEF-H1 activates RhoA by exchanging GDP for GTP, allowing activation of the downstream kinase ROCK. ROCK-dependent phosphorylation of myosin light chain (MLC) will then induce actin stress fibers formation.

Interestingly, BepC and GEF-H1 were found to form a trimeric complex together with MRCKα, however, it remained unclear whether this kinase also interacts with BepC, or whether it binds primarily via GEF-H1 to which it was also found to bind in the absence of BepC. Yet, we can conclude that the participation of MRCKα is neglectable for BepC-triggered cytoskeletal changes given that a full knock-out of MRCKα did not interfere with actin stress fiber formation mediated by the effector. Nevertheless, it is also conceivable that under relevant physiological conditions prone to activation of MRCKα, the interaction with GEF-H1 may contribute to BepC-triggered stress fiber formation via direct phosphorylation of MLC and inhibition of myosin light chain phosphatase (MLCP) [42] (see model in Fig. 4A).

The high level of sequence conservation between BepC homologs [16] and the consistency in the ability to trigger actin rearrangements indicate an evolutionary conserved molecular function that is playing a major role in the context of a shared infection strategy of the bartonellae [6, 7]. Thus, future work should place this effector signaling mechanism into a larger pathophysiological context of *Bartonella* spp. infection in the established infection models for invasome formation and alternative modes of bacterial internalization [12, 24-26], migration of infected dendritic cells [11], and related innate immune cell functions [5, 43].

## Material and Methods

### Bacterial strains and growth conditions

The bacterial strains used in this study are listed in Table S2. *Bartonella* species were grown on Columbia blood agar (CBA, Oxoid, CM0331) plates containing 5% defibrinated sheep blood (Oxoid, SR0051) at 35°C and 5% CO_2_ for 3 days then expended for 2 days on new plates. When necessary, media were supplemented with 30 μg/ml kanamycin, 100 μg/ml streptomycin. *E. coli* strain was cultivated in Luria-Bertani liquid medium (LB) or on LB agar on plates (LA) at 37°C overnight. Media were supplemented with 50 μg/ml kanamycin and 1 mM diaminopimelic acid (DAP, Sigma, D1377).

### Construction of plasmids used in this work

*Bartonella* expression plasmids used in this study are listed in Table S3. Eukaryotic expression plasmids used in this study are listed in Table S4. Plasmids construction details as summarized in Table S5 and the used PCR primers as listed in Table S6.

### Conjugation of *Bartonella*-expression plasmids into *Bartonella*

*Bartonella henselae* Δ*bepA-G* (MSE150) was grown on CBA plates in presence of 100 μg/ml streptomycin at 35°C and 5% CO_2_ for 3 days then expanded on new plates for 2 days. The day before conjugation, 5 ml of LB containing 1 mM DAP and 50 μg/ml kanamycin were inoculated with a conjugation strain (JKE170) containing the plasmid of interest. After overnight incubation at 37°C, a subculture was prepared by inoculating 5 ml of LB containing 1 mM DAP and 50 μg/ml kanamycin with 200 µl of overnight culture before being incubated for 2 h at 37°C. In order to remove antibiotics, 500 µl of the subculture was centrifuged for 4 min at 2’000 x g and the bacterial pellet was resuspended in 500 µl of M199 (Gibco, 22340-020) supplemented with 10% of heat-inactivated fetal calf serum (FCS), the washing step was repeated once. The same process was applied to *Bartonella*, bacteria were harvested in 1 ml of M199 10% heat-inactivated FCS and centrifuged for 4 min at 2’000 x g. The bacterial pellet was resuspended in 500 µl of M199 10% heat-inactivated FCS before being centrifuged again and resuspended in 100 µl of M199 10% heat-inactivated FCS. 20 µl of *E. coli* were mixed with 100 µl of *Bartonella* and incubated for 5 h at 35°C, 5% CO_2_ on a nitrocellulose filter deposited on a CBA plate supplemented with 1 mM of DAP. The filter was transferred in an Eppendorf tube containing 1 ml of M199 10% heat-inactivated FCS and the bacteria were resuspended by gently shaking. 50 µl were plated on a CBA plate supplemented with 30 μg/ml kanamycin and 100 μg/ml streptomycin. A single colony was selected and subsequently tested to confirm the presence of the plasmid.

### Cell culture

Human umbilical vein endothelial cells (HUVEC) were isolated as described before (Dehio et al., 1997) and cultured at 37°C with 5% CO_2_ in Endothelial Cell Growth Medium (ECGM, Promocell, C-22010) supplemented with Endothelial Cell Growth Medium SupplementMix (Promocell, C-39215). HeLa cells were cultured at 37°C with 5% CO_2_ in DMEM (Sigma, D6429) supplemented with 10% heat-inactivated FCS.

### Cell infection for microscopy

HUVECs were plated at a density of 3’000 cells/well in a 96-well plate (Corning, #3904) pre-coated with 0.2% of gelatin using supplemented ECGM. HeLa cells were seeded at a density of 12’500 cells/well in a µ-Slide 8 Well (Ibidi, Cat. N°:80826) or at a density of 2’000 cells/well in a 96-well plate (Corning, #3904) using DMEM supplemented with 10% heat-inactivated FCS. The next day, cells were infected with *Bartonella* at the indicated MOI in M199 (Gibco, 22340-020) supplemented with 10% of heat-inactivated FCS in presence of 10 µM of isopropyl-β-D-thiogalactoside (IPTG, Biochemica, A1008). After incubation at 35°C and 5% CO_2_, cells were fixed with 3.7% of paraformaldehyde for 10 minutes and washed 3 times with PBS.

### Cell transfection

HeLa cells were seeded at a density of 12’500 cells/well in a µ-Slide 8 Well (Ibidi, Cat. N°:80826) or at a density of 2’000 cells/well on a 96-well plate (Corning, #3904) using DMEM supplemented with 10% of heat-inactivated FCS. The next day, cells were transfected according to manufacturer instruction with a transfection mix containing a ratio of 1 µg of plasmid for 2 µl FuGene HD transfection reagent (Promega, REF E2311) diluted in DMEM without FCS. After transfection for 24 h at 37°C and 5% CO_2_, cells were fixed with 3.7% of paraformaldehyde for 10 minutes and washed 3 times with PBS.

### Immunostaining

Fixed cells were permeabilized for 10 minutes with PBS 0.2% BSA (Sigma, A9647) and 0.5% Triton X-100 (Sigma, T9284). After being washed 3 times with PBS with 0.2% BSA, cells were incubated overnight at 4°C in the presence of the primary antibody (Table S7) diluted in PBS with 0.2% BSA. After 2 more washes with PBS with 0.2% BSA, cells were incubated for 2 h in the dark in presence of the secondary antibody (Table S7), DAPI (Sigma, D9542, 1 µg/ml) and, when indicated, DY-547P1 phalloidin (Dyomics GmbH, final concentration 1/250) diluted in PBS with 0.2% BSA. Cells were finally washed 3 times with PBS. 96-well plates were imaged with an MD ImagXpress Micro automated microscope from Molecular devices and fluorescence was detected at 10x magnification. Images were processed in MetaXpress. For µ-Slide, the stained samples were analyzed using a LEICA point scanning confocal “SP8” microscope (Imaging Core Facility, Biozentrum, University of Basel, Switzerland). Z-stacks with 34-40 focal planes with a spacing of 0.45 µm were recorded and images were reconstructed by Z-projection using ImageJ.

### Pull-down assay

HeLa cells or HUVECs were plated in round plates (Falcon, REF 353003) at a density of 365’000 or 544’000 cells per plate, respectively. The cells were then incubated overnight at 37°C with 5% CO_2_ in DMEM complemented with 10% heat-inactivated FCS. In the morning, cells have been infected with the indicated strain of *Bartonella* at MOI of 200 for 24 h at 35°C with 5% CO_2_ in M199 supplemented with 10% heat-inactivated FCS in presence of 10 µM of IPTG. After infection, cells were washed 3 times with ice-cold PBS and lysed with lysis buffer containing 50 mM Hepes (pH 7.5), 150 mM NaCl, protease inhibitor (Roche, 11836170001), and 1% Nonidet P40 substitute (Sigma, 74385). Cell lysates were collected with a cell scraper and incubated 30 minutes on ice. After centrifugation at 20’000 x g for 30 min at 4°C, the supernatants were incubated in presence of 20 µl of protein G agarose beads (Roche, 11243233001) for 3 h at 4°C on a rotor to reduce unspecific binding. After removing the beads by centrifugation for 30 seconds at 12’000 x g, 2 µg of antibody (Table S7) was added to the supernatant. After 3 h of incubation at 4°C on a rotor, 20 µl of protein G agarose was added to the lysates and incubated overnight at 4°C on a rotor. The next morning, agarose beads were collected by centrifugation for 30 seconds at 12’000 x g before being washed 2 times with lysis buffer and 2 more times with lysis buffer without NP-40. Proteins were eluted from the beads by incubation at 95°C for 10 minutes in SDS sample buffer. Elution fractions and cell lysates before pull-down were analyzed by immunoblot. The same protocol was applied for samples analyzed by mass spectrometry although one cell culture flask of 150 cm2 was used per infection and that proteins were eluted by incubation at 95°C for 10 minutes with 2% SDS.

### Sample preparation for mass spectrometry

Proteins were precipitated with trichloroacetic acid and incubated 10 min at 4°C. The protein pellet was washed twice with cold acetone and resuspended with 4 M urea. Then the samples were treated with 5 mM of tris(2-carboxyethyl)phosphine (TCEP) for 30 min at 37°C in order to reduce disulfide bonds. After incubation, iodoacetamide (1.8 mg/ml final) was added to the samples to irreversibly prevent the formation of disulfide bonds and incubated for 30 min at 25°C in the dark. The samples were subsequently diluted with 0.1 M ammonium bicarbonate to have a final concentration of urea of 1.6 M. For digestion, the proteins were incubated overnight at 37°C in presence of 1 µg of trypsin. After acidification with trifluoroacetic acid (TFA, 1% final), the peptides were loaded on a C-18 column (The Nest Group, SS18V) pre-equilibrated with buffer A (0.1% TFA). The column was washed 3 times with buffer C (5% acetonitrile / 95%water (v/v) and 0.1% TFA) and peptides were eluted with buffer B (50% acetonitrile / 50%water (v/v) and 0.1% TFA). The peptides were finally dried under vacuum and kept at - 80°C. Before LC-MS/MS mass analysis, samples were resuspended in 0.1% formic acid by sonication.

### Mass Spectrometry Analysis

For each sample, aliquots of 0.4 μg of total peptides were subjected to LC-MS analysis using a dual pressure LTQ-Orbitrap Elite mass spectrometer connected to an electrospray ion source (both Thermo Fisher Scientific) and a custom-made column heater set to 60°C. Peptide separation was carried out using an EASY nLC-1000 system (Thermo Fisher Scientific) equipped with a RP-HPLC column (75μm × 30cm) packed in-house with C18 resin (ReproSil-Pur C18–AQ, 1.9 μm resin; Dr. Maisch GmbH, Germany) using a linear gradient from 95% solvent A (0.1% formic acid in water) and 5% solvent B (80% acetonitrile, 0.1% formic acid, in water) to 35% solvent B over 50 minutes to 50% solvent B over 10 minutes to 95% solvent B over 2 minutes and 95% solvent B over 18 minutes at a flow rate of 0.2 μl/min. The data acquisition mode was set to obtain one high resolution MS scan in the FT part of the mass spectrometer at a resolution of 120,000 full width at half maximum (at 400 m/z, MS1) followed by MS/MS (MS2) scans in the linear ion trap of the 20 most intense MS signals. The charged state screening modus was enabled to exclude unassigned and singly charged ions and the dynamic exclusion duration was set to 30 s. The collision energy was set to 35%, and one microscan was acquired for each spectrum.

### Protein identification and label-free quantification

The acquired raw-files were imported into the Progenesis QI software (v2.0, Nonlinear Dynamics Limited), which was used to extract peptide precursor ion intensities across all samples applying the default parameters. The generated mgf files were searched using MASCOT against a decoy database containing normal and reverse sequences of the concatenated Homo sapiens (UniProt, Mai 2016) and Bartonella henselae (UniProt, July 2016) proteome and commonly observed contaminants (in total 44102 sequences) generated using the SequenceReverser tool from the MaxQuant software (Version 1.0.13.13). The following search criteria were used: full tryptic specificity was required (cleavage after lysine or arginine residues, unless followed by proline); 3 missed cleavages were allowed; carbamidomethylation (C) was set as fixed modification; oxidation (M) and protein N-terminal acetylation were applied as variable modifications; mass tolerance of 10 ppm (precursor) and 0.6 Da (fragments) was set. The database search results were filtered using the ion score to set the false discovery rate (FDR) to 1% on the peptide and protein level, respectively, based on the number of reverse protein sequence hits in the datasets. Quantitative analysis results from label-free quantification were normalized and statically analyzed using the SafeQuant R package v.2.3.4 (https://github.com/eahrne/SafeQuant/) (PMID: 27345528) to obtain protein relative abundances. This analysis included summation of peak areas per protein and LC MS/MS run followed by calculation of protein abundance ratios. Only isoform specific peptide ion signals were considered for quantification. The summarized protein expression values were used for statistical testing of differentially abundant proteins between conditions. Here, empirical Bayes moderated t-Tests were applied, as implemented in the R/Bioconductor limma package (http://bioconductor.org/packages/release/bioc/html/limma.html). The resulting p-values were adjusted for multiple testing using the Benjamini Hochberg method.

All LC-MS analysis runs are acquired from independent biological samples. To meet additional assumptions (normality and homoscedasticity) underlying the use of linear regression models and Student t-Test MS-intensity signals are transformed from the linear to the log-scale.

Unless stated otherwise linear regression was performed using the ordinary least square (OLS) method as implemented in base package of R v.3.1.2 (http://www.R-project.org/). The sample size of three biological replicates was chosen assuming a within-group MS-signal Coefficient of Variation of 10%. When applying a two-sample, two-sided Student’s t-test this gives adequate power (80%) to detect protein abundance fold changes higher than 1.65, per statistical test. Note that the statistical package used to assess protein abundance changes, SafeQuant, employs a moderated t-Test, which has been shown to provide higher power than the Student’s t-test. We did not do any simulations to assess power, upon correction for multiple testing (Benjamini-Hochberg correction), as a function of different effect sizes and assumed proportions of differentially abundant proteins.

### Inhibitor treatment of infected HeLa cells

HeLa cells were seeded at a density of 2’000 cells/well on a 96-well plate (Corning, #3904) using DMEM supplemented with 10% of heat-inactivated FCS. After overnight incubation at 37°C with 5% CO_2_, cells were infected with *Bhe* Δ*bepA-G* carrying the empty plasmid or expressing BepC at MOI of 200 in M199 (Gibco, 22340-020) supplemented with 10% of heat-inactivated FCS in presence of 10 µM of IPTG. After 24 h of incubation at 35°C and 5% CO_2_, the medium was removed and cells were incubated with inhibitor diluted in DMEM at 35°C with 5% CO_2_. The treatment consisted of 2 µg/ml of Rho inhibitor I (Cytoskeleton, CT04) for 2 h or 20 µM of Y27632 (Sigma, Y0503) for 1 hour. The experiment was stopped by fixation with 3.7% of paraformaldehyde for 10 minutes. Finally, the cells were washed 3 times with PBS before being stained and imaged by microscopy.

### Generation of knock-out cells

HeLa cells were seeded at a density of 1’400’000 cells per 150 cm2 flask in DMEM supplemented with 10% of heat-inactivated FCS and incubated overnight at 37°C with 5% CO_2_. The day after, in 1.2 ml of DMEM without FCS, 12 µg of plasmid encoding GFP and the guide RNA (gRNA) targeting the first exon of the gene of interest were mixed with 12 µg of plasmid carrying the puromycin resistance gene and the gRNA targeting the last exon. After adding 48 µl of FuGene HD transfection reagent (Promega, E2311), the transfection mix was incubated for 15 min at room temperature before being transferred in the cell culture flask. The cells were transfected for 24 h at 37 °C with 5% CO_2_. Double-transfected cells were selected in the presence of puromycin (1.5 µg/ml) for 24 h at 37°C with 5% CO_2_ followed by FACS to select GFP-positive cells. Selected cells were collected in DMEM with 10% heat-inactivated FCS supplemented with penicillin-streptomycin and expanded for several days. Finally, cells were stored at −80°C in DMEM supplemented with 10% heat-inactivated FCS and 10% DMSO. The expression level of the protein of interest was monitored via immunoblot.

### Immunoblot analysis

The samples used for immunoblot analysis were separated by SDS-PAGE on a 4-20% gradient gel (Mini-PROTEAN TGX Gels, Biorad, Cat# 456-1093). Gel electrophoresis was performed at 120 V in running buffer (Tris-glycine, 0.1% SDS). Proteins were transferred on a PVDF membrane (GE Healthcare, 10600021) via wet electroblotting at 100 V in transfer buffer (20% methanol, Tris-glycine) at 4°C. After transfer, the membrane was incubated for 1 hour in blocking buffer (PBS, 0.1% Tween 20 (Sigma, 93773), supplemented with 5% milk or 5% BSA according to antibody recommendation). After washing with PBS 0.1% Tween 20, the membrane was incubated overnight at 4°C in blocking buffer with the primary antibody (Table S7). The membrane was washed again with PBS 0.1% Tween 20 before being incubated 1 hour at room temperature in blocking buffer with the secondary antibody (Table S7). The blots were developed using LumiGLO Reserve Chemiluminescent Substrate System (KPL, 54-70-00, 54-69-00). Finally, the signal was detected with LAS4000 (Fujifilm).

### Quantification of pMLC via Cellprofiler

Experiments performed in 96-well plates were subjected to automated microscopy, using MD ImageXpress Micro automated microscopes. For each condition, at least 6 wells with 25 sites were imaged in 4 different wavelengths corresponding to the applied cell staining (DAPI, DY-547P1 phalloidin, pMLC, *Bartonella*). Images were analyzed with the CellProfiler software [38]. Two separate Cellprofiler pipelines are used for each assay. The first pipeline calculates a shading model, which is used by the second pipeline to correct images prior to analysis. To correct uneven illumination inherent in wide-field microscopic imaging (shading), an illumination function was computed. The illumination function was calculated on all images based on the Background method. The resulting image was smoothed using a Gaussian method with a 100-filter size. To reduce the signal originating from the bacterial DNA in the DAPI channel, the signal corresponding to *Bartonella* was subtracted from the DAPI image. On all images, CellProfiler was executed to perform object segmentation and measurements with the following steps. Nuclei were detected as primary objects using an Automatic strategy and clumped objects were identified based on their shape and segmented based on their intensity. HeLa cells were detected as secondary objects via their DY-547P1 phalloidin signal by using a Propagation method from the nuclei followed by a Global threshold strategy combined with an Otsu threshold method. The average of the mean intensity of each cell within one site was measured for pMLC signal. Data from all sites from the same conditions were compiled together. The mean cell intensity per site was normalized on the uninfected condition.

### Statistical analysis

Graph was generated with GraphPad Prism 8. Statistical analyses were performed using Kruskal-Wallis test with Dunn’s multiple comparison test. For the graph presented in the figures, significance was denoted P∗∗∗∗ < 0.0001.

### Software

ImageJ [39] was used to create z-projection and x-z sections of confocal microscopy images. MetaXpress (Molecular Devices) was used to acquire and generate microscopy pictures from automated microscope. Image analysis and the calculation of the average of the mean cell pMLC fluorescence intensity was realized via Cellprofiler [38]. GraphPad prism 8 (GraphPad) was used to create the dot-plot and the statistical analysis of pMLC phosphorylation. Geneious Prime 2019 (Geneious) was used to design cloning of plasmids. The schematic model of BepC-mediated actin stress fiber formation was created with Biorender.com.

## Acknowledgments

We thank Sarah Stiegeler and Drs. Raquel Conde-Alvarez, Simone Eicher and Kathrin Pieles for sharing initial findings on the BepC phenotype and BepC/GEF-H1 interaction, and Drs. Maxime Québatte and Alexander Harms for sharing an unpublished *Bartonella* expression vector. We are grateful to Drs. Isabel Sorg and Maxime Québatte for critically reading of the manuscript and helpful comments. We thank the Proteomics Core Facility for performing mass spectrometry analyses, the FACS Core Facility for cell sorting, and the Imaging Core Facility for providing technical support. We also thank Prof. Perihan Nalbant (University of Duisburg-Essen, Germany) for providing the eGFP-GEF-H1 mammalian expression vector. The plasmids used to generate CRISPR/Cas knock-out cell lines, pSpCas9(BB)-2A-GFP (PX458) (Addgene plasmid # 48138; http://n2t.net/addgene:48138; RRID:Addgene_48138) and pSpCas9(BB)-2A-Puro (PX459) (Addgene plasmid # 48139; http://n2t.net/addgene:48139; RRID:Addgene_48139), were a gift from Feng Zhang. The Bruderholzspital Basel is acknowledged for providing human umbilical cords.

## Autor Contributions

Conceived and designed the experiments: SM CD. Performed the experiments: SM. Analyzed the data: SM. Contributed reagents/materials/analysis tools: SM. Wrote the paper: SM CD.

## Supplementary Information

**Fig. S1.**
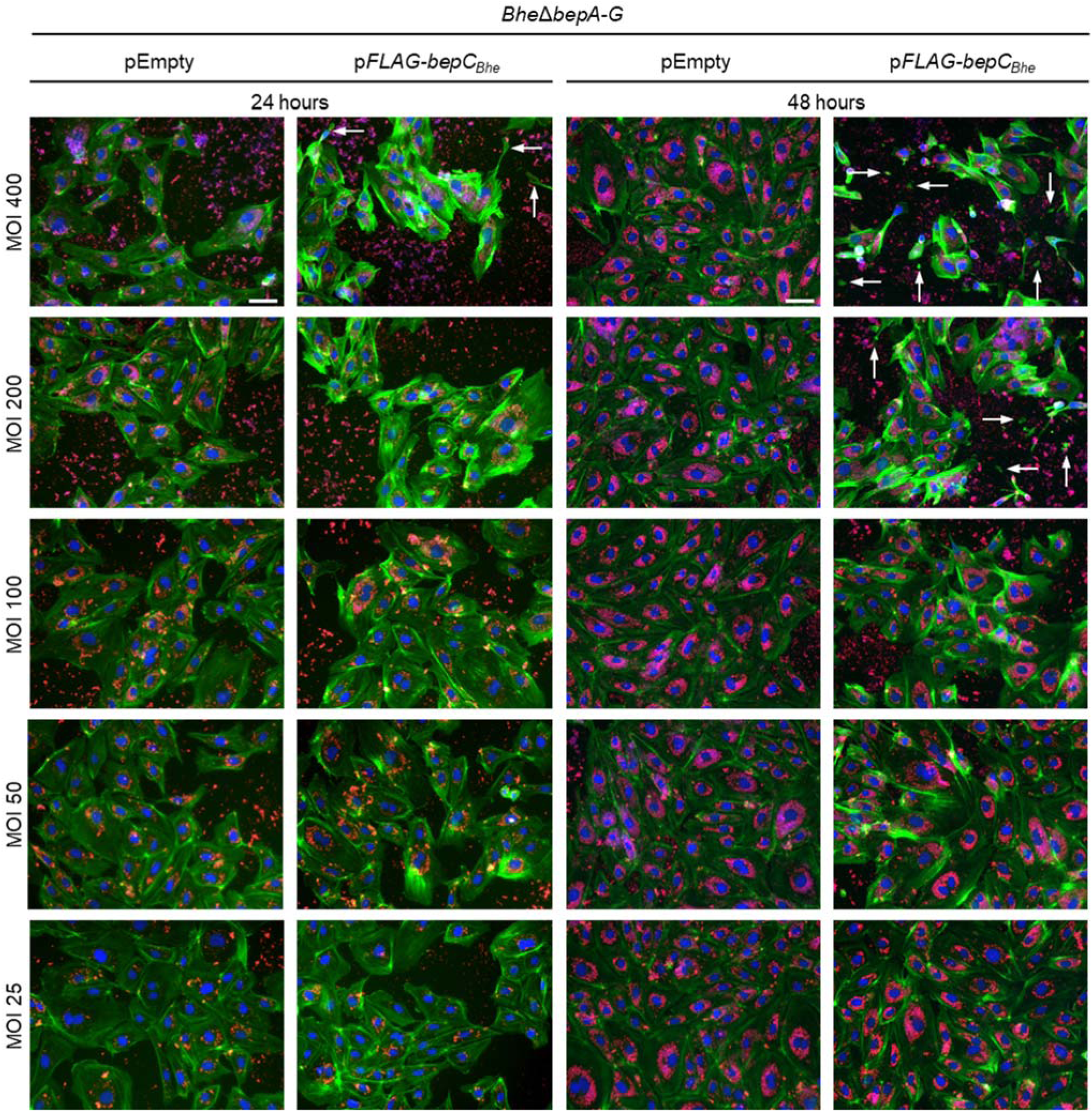
BepC_*Bhe*_-triggered actin stress fiber formation in infected *B. henselae-*infected HUVEC is dependent on time and multiplicity of infection. HUVECs were infected with *Bhe* Δ*bepA-G* expressing FLAG-tagged BepC_*Bhe*_ or carrying empty plasmid as a negative control at the indicated MOIs for 24 or 48 h. After fixation, cells were stained by immunocytochemistry, followed by fluorescence microscopy analysis. F-actin is represented in green, DNA in blue, and bacteria in red (scale bar = 50 µm). Arrows indicate cell fragments as a result of the cell fragmentation phenotype.

**Fig. S2.**
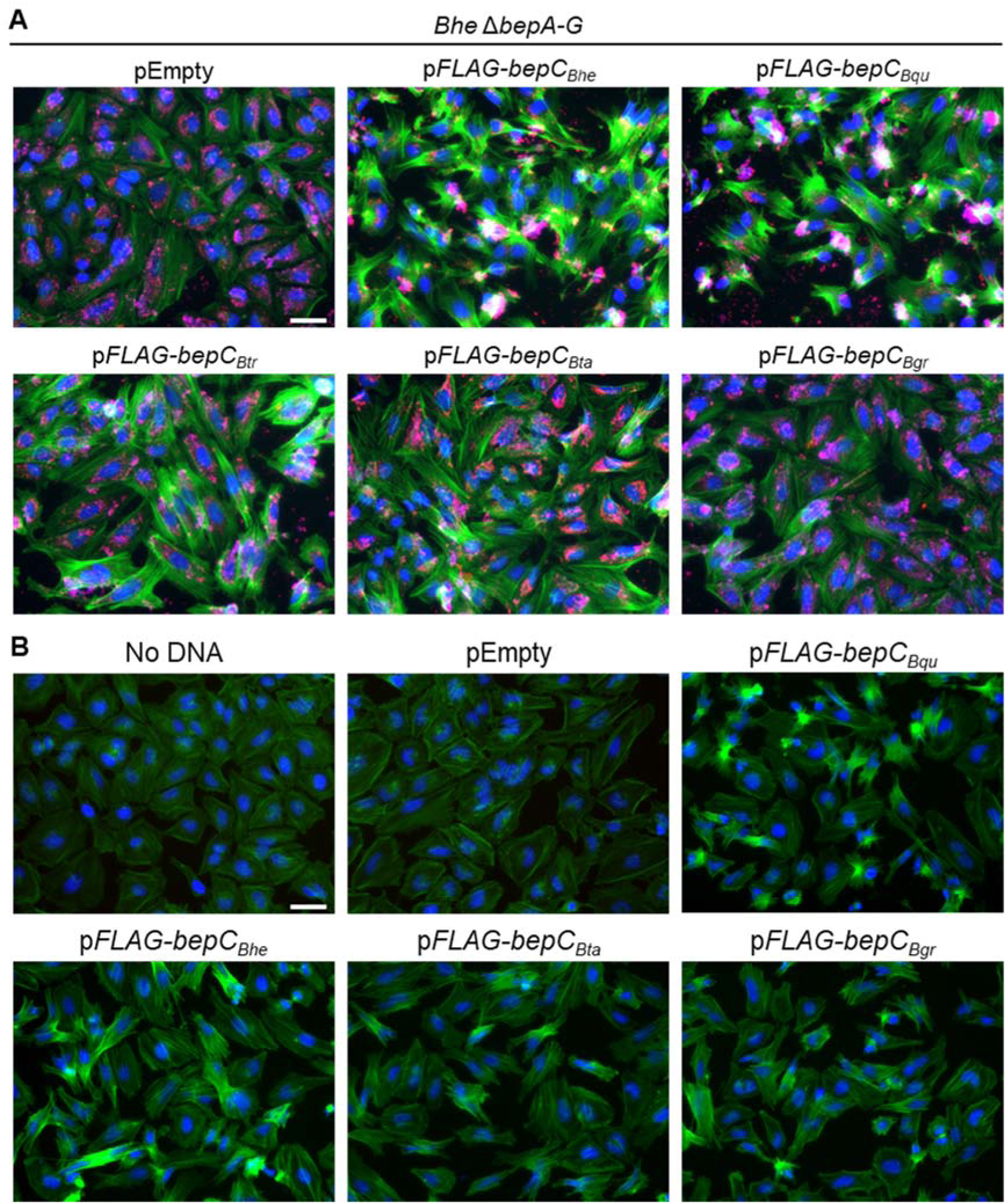
BepC-triggered actin stress fiber formation is conserved among homologs encoded by various *Bartonella* species. (**A**) HeLa cells were infected with the indicated isogenic *Bhe* Δ*bepA-G* expressing BepC homologs at MOI of 400. (**B**) HeLa cells were transfected with indicated expression plasmids encoding different BepC homologs. After 24 h of transfection or 48 h of infection, cells were fixed and immunocytochemically stained, followed by fluorescence microscopy analysis. F-actin is represented in green, DNA in blue, and bacteria in red (scale bar = 50 µm). *Bhe* (*B. henselae*); *Bqu* (*B. quintana*); *Btr* (*B. tribocorum*); *Bta* (*B. taylorii*); *Bgr* (*B. grahamii*).

**Fig S3.**
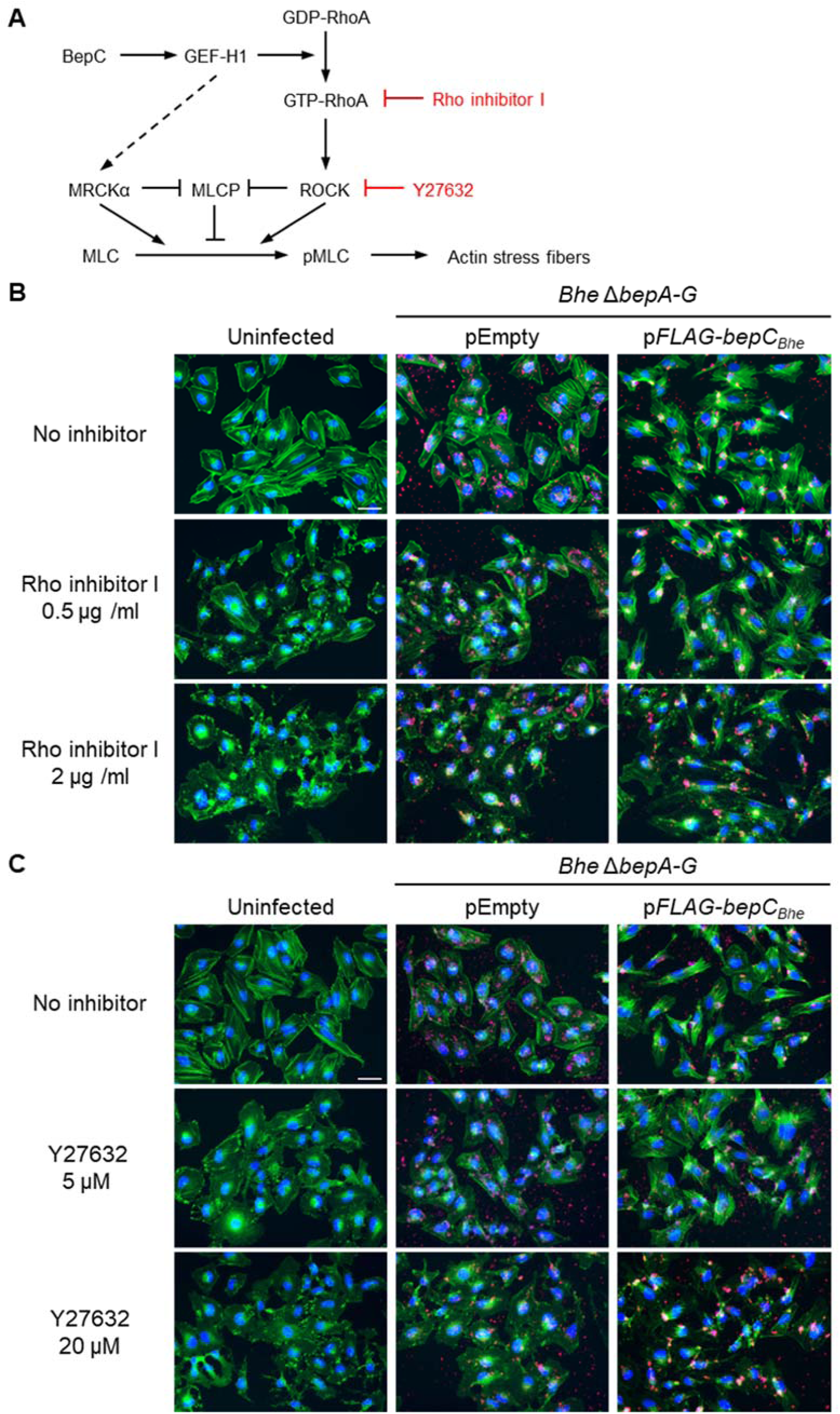
Inhibition of RhoA/B/C or ROCK reduces actin stress fiber formation mediated by BepC_*Bhe*_. (**A**) Proposed model of BepC-triggered actin stress fiber formation via the activation of the RhoA pathway and the targets of inhibitors used for validation. (**B-C**) HeLa cells were infected with *Bhe* Δ*bepA-G* expressing FLAG-tagged BepC_*Bhe*_ or carrying the empty plasmid as a negative control at MOI of 200. After 24 h of infection, cells were treated with inhibitors as specified below, followed by fixation and immunocytochemical staining. Specimen were then analyzed by fluorescence microscopy. F-actin is represented in green, DNA in blue, and bacteria in red (scale bar = 50 µm). (**B**) HeLa cells incubated for 2 h in the presence or absence of Rho inhibitor I. (**C**) HeLa cells incubated for 1 h in the presence or absence of the ROCK inhibitor Y27632.

**Fig. S4.**
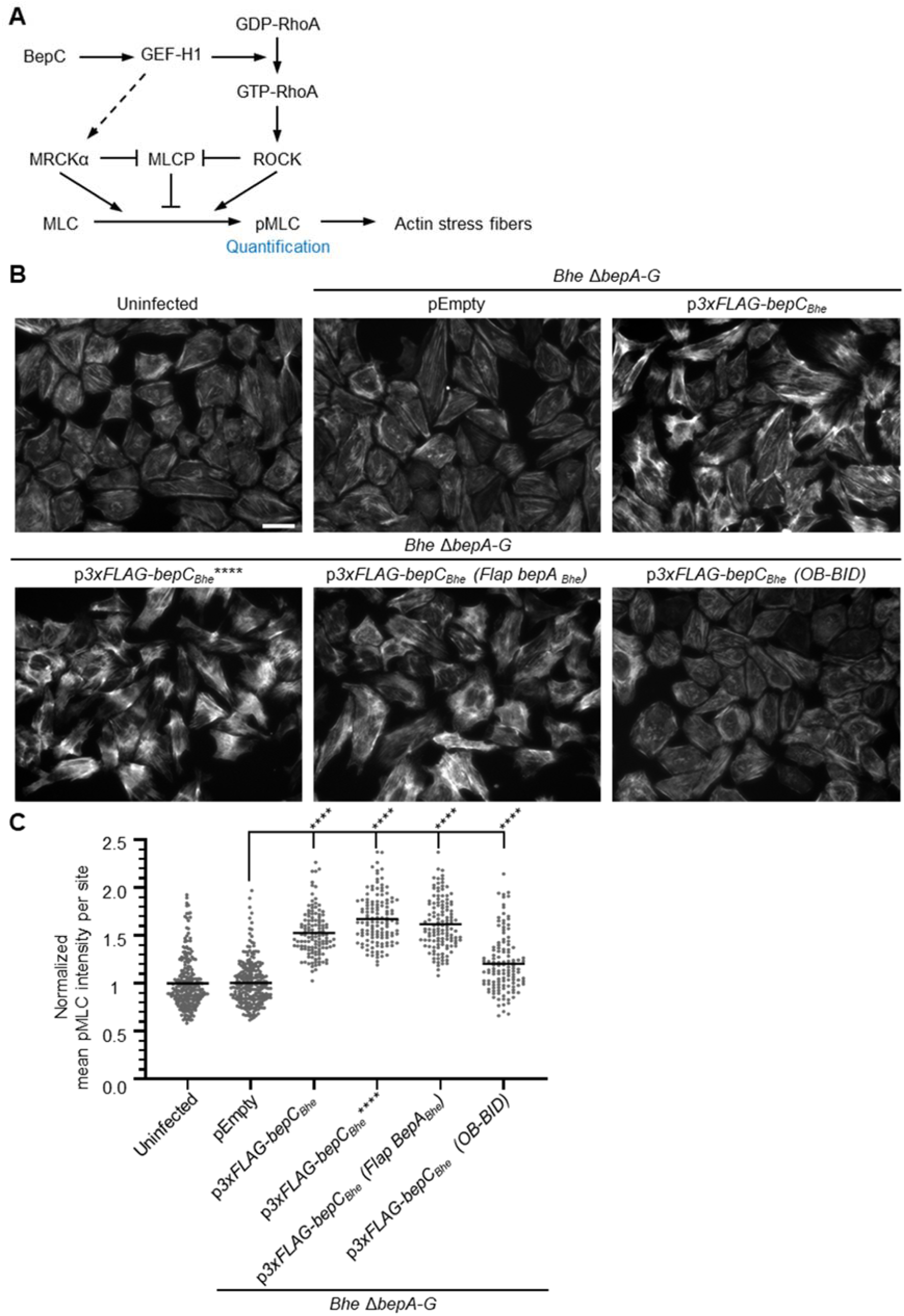
BepC_*Bhe*_ induces a significant increase of myosin light chain phosphorylation. (**A**) Proposed model of BepC-triggered actin stress fiber formation. (**B**) HeLa cells were infected with isogenic *Bhe* Δ*bepA-G* strains expressing FLAG-tagged BepC_*Bhe*_ wild-type or mutant variants, or carrying the empty plasmid at multiplicity of infection (MOI) of 200. After 48 h of infection, cells were fixed and immunocytochemically stained, followed by fluorescence microscopy analysis. Phosphorylated myosin light chain (pMLC) is represented in white (scale bar = 50 µm). BepC_*Bhe*_**** = BepC_*Bhe*_ H146A, K150A, R154A, R157A. (**C**) HeLa cells infected as described in (B) were fixed and immunocytochemically stained for DNA, bacteria, F-actin and phosphorylated myosin light chain (pMLC) and imaged by fluorescence microscopy. The mean fluorescence intensity of pMLC was quantified for each individual cell using CellProfiler. Data are represented as dot plots with each data point corresponding to the average of all mean cell intensity values within one imaged site. Statistical significance was determined using Kruskal-Wallis test (**** corresponds to p-value ≤ 0.0001).

**Table S1.** List of proteins identified by Mass spectrometry (see Fig. 3.A for details of the approach).

*Table S1 was submitted separately from the combined pdf file of the manuscript and supplementary information.*

**Supplementary Table S2.** List of bacterial strains used in this study

| Strain name | Description | Reference |
| --- | --- | --- |
| MSE150 | *Bhe ΔbepA-G* | [8] |
| SIM B1-45 | *Bhe ΔbepA-G* / pFLAG-*bepC<sub>Bgr</sub>* (MSE150 containing pSIM051) | This work |
| SIM B1-46 | *Bhe ΔbepA-G* / pFLAG-*bepC<sub>Bqu</sub>* (MSE150 containing pSIM054) | This work |
| SIM B1-49 | *Bhe ΔbepA-G* / pFLAG-*bepC<sub>Bta</sub>* (MSE150 containing pSIM058) | This work |
| SIM B1-52 | *Bhe ΔbepA-G* / pEmpty (MSE150 containing pBZ485_a_empty) | This work |
| SIM B2-06 | *Bhe ΔbepA-G* / pFLAG-*bepC<sub>Bhe</sub>* (MSE150 containing pSIM091) | This work |
| SIM B2-07 | *Bhe ΔbepA-G* / pFLAG-*bepC<sub>Btr</sub>* (MSE150 containing pSIM062) | This work |
| SIM B2-08 | *Bhe ΔbepA-G* / pFLAG- *bepC<sub>Bhe</sub>* **** (MSE150 containing pSIM081) | This work |
| SIM B2-16 | *Bhe ΔbepA-G* / pFLAG-*bepC<sub>Bhe</sub>* (*OB-BID*) (MSE150 containing pSIM098) | This work |
| SIM B2-25 | *Bhe ΔbepA-G* / pFLAG-*bepC<sub>Bhe</sub>* (*Flap bepA<sub>Bhe</sub>*) (MSE150 containing pSIM117) | This work |
| SIM B2-18 | *Bhe ΔbepA-G* / p3xFLAG-*bepC<sub>Bhe</sub>* (MSE150 containing pSIM107) | This work |
| SIM B2-38 | *Bhe ΔbepA-G* / p3xFLAG- *bepC<sub>Bhe</sub>* **** (MSE150 containing pSIM127) | This work |
| SIM B2-42 | *Bhe ΔbepA-G* / p3xFLAG-*bepC<sub>Bhe</sub>* (*OB-BID*) (MSE150 containing pSIM131) | This work |
| SIM B2-43 | *Bhe ΔbepA-G* / p3xFLAG-*bepC<sub>Bhe</sub>* (*Flap bepA<sub>Bhe</sub>*) (MSE150 containing pSIM132) | This work |
| JKE170 | *E.coli* strain used to conjugate *Bartonella* expression vectors in *Bhe ΔbepA-G* | [40] |

**Supplementary Table S3.** List of *Bartonella* expression vectors used in this work

| Internal name | Plasmid name | Description | Reference |
| --- | --- | --- | --- |
| pCD353 |  | *Bartonella* spp. vector encoding GFP | [41] |
| pBZ485_a_empty | pEmpty | pSIM037 digested with NdeI, empty vector | This work |
| pSIM037 | pBZ485_a_GFP | Derivative of pCD353 encoding GFP | This work |
| pSIM051 | pFLAG-bepC<sub>Bgr</sub> | pBZ485_a_empty encoding FLAG-BepC<sub>Bgr</sub> | This work |
| pSIM054 | pFLAG-bepC<sub>Bqu</sub> | pBZ485_a_empty encoding FLAG-BepC<sub>Bqu</sub> | This work |
| pSIM058 | pFLAG-bepC<sub>Bta</sub> | pBZ485_a_empty encoding FLAG-BepC<sub>Bta</sub> | This work |
| pSIM062 | pFLAG-bepC<sub>Btr</sub> | pBZ485_a_empty encoding FLAG-BepC<sub>Btr</sub> | This work |
| pSIM081 | pFLAG-bepC<sub>Bhe</sub> **** | pBZ485_a_empty encoding FLAG-BepC<sub>Bhe</sub> **** | This work |
| pSIM091 | pFLAG-bepC<sub>Bhe</sub> | pBZ485_a_empty encoding FLAG-BepC<sub>Bhe</sub> | This work |
| pSIM098 | pFLAG-bepC<sub>Bhe</sub> (OB-BID) | pBZ485_a_empty encoding FLAG-BepC<sub>Bhe</sub> (OB-BID) | This work |
| pSIM107 | p3xFLAG-bepC<sub>Bhe</sub> | pBZ485_a_empty encoding 3xFLAG-BepC<sub>Bhe</sub> | This work |
| pSIM117 | pFLAG-bepC<sub>Bhe</sub> (Flap BepA<sub>Bhe</sub>) | pBZ485_a_empty encoding FLAG-BepC<sub>Bhe</sub> (Flap BepA<sub>Bhe</sub>) | This work |
| pSIM127 | p3xFLAG-bepC<sub>Bhe</sub> **** | pBZ485_a_empty encoding 3xFLAG-BepC<sub>Bhe</sub> **** | This work |
| pSIM131 | p3xFLAG-bepC<sub>Bhe</sub> (OB-BID) | pBZ485_a_empty encoding 3xFLAG-BepC<sub>Bhe</sub> (OB-BID) | This work |
| pSIM132 | p3xFLAG-bepC<sub>Bhe</sub> (Flap BepA<sub>Bhe</sub>) | pBZ485_a_empty encoding 3xFLAG-BepC<sub>Bhe</sub> (Flap BepA<sub>Bhe</sub>) | This work |

**Supplementary Table S4.** List of eukaryotic expression plasmids used in this work

| Internal name | Plasmid name | Description | Reference |
| --- | --- | --- | --- |
| pFlag-CMV2 | pEmpty | eukaryotic expression vector encoding a short FLAG tag | Eastman Kodak, New Haven |
| pCMV5-eGFP-GEF-H1 | *peGFP-GEF-H1* | eukaryotic expression vector encoding eGFP-GEF-H1 | Obtained from P. Nalbant - [32] |
| pX458 | pSpCas9(BB)-2A-GFP | eukaryotic expression vector encoding Cas9 and GFP | [42] |
| pX459 | pSpCas9(BB)-2A-Puro | eukaryotic expression vector encoding Cas9, Puro<sup>R</sup> | [42] |
| pSIM138 |  | pX458 with sgRNA targeting Exon1 of GEF-H1 | This work |
| pSIM139 |  | pX459 with sgRNA targeting Exon22 of GEF-H2 | This work |
| pSIM140 |  | pX458 with sgRNA targeting Exon1 of MRCKa | This work |
| pSIM141 |  | pX459 with sgRNA targeting Exon36 of MRCKa | This work |
| pSIM142 | *pFLAG-bepC<sub>Bgr</sub>* | pFlag-CMV2 encoding FLAG-BepC<sub>Bgr</sub> | This work |
| pSIM143 | *pFLAG-bepC<sub>Bqu</sub>* | pFlag-CMV2 encoding FLAG-BepC<sub>Bqu</sub> | This work |
| pSIM144 | *pFLAG-bepC<sub>Bta</sub>* | pFlag-CMV2 encoding FLAG-BepC<sub>Bta</sub> | This work |
| pSIM146 | *pFLAG-bepC<sub>Bhe</sub>* | pFlag-CMV2 encoding FLAG-BepC<sub>Bhe</sub> | This work |
| pSIM147 | *p3xFLAG-bepC<sub>Bhe</sub>* | pFlag-CMV2 encoding 3xFLAG-BepC<sub>Bhe</sub> | This work |
| pSIM148 | *p3xFLAG-bepC<sub>Bhe</sub> (FIC-OB)* | pFlag-CMV2 encoding 3xFLAG-BepC<sub>Bhe</sub> (FIC-OB) | This work |
| pSIM150 | *p3xFLAG-bepC<sub>Bhe</sub> \*\*\*\** | pFlag-CMV2 encoding 3xFLAG-BepC<sub>Bhe</sub> **** | This work |
| pSIM154 | *p3xFLAG-bepC<sub>Bhe</sub> (OB-BID)* | pFlag-CMV2 encoding 3xFLAG-BepC<sub>Bhe</sub> (OB-BID) | This work |

**Supplementary Table S5.** Construction details for plasmids used in this work

| Plasmid | Insert template | PCR 1 | PCR2 | Backbone | Restriction |
| --- | --- | --- | --- | --- | --- |
| pSIM051 | Genomic DNA | prSIM106/prSIM110 | prSIM076/prSIM110 | pSIM037 | SacI/NotI |
| pSIM054 | Genomic DNA | prSIM107/prSIM111 | prSIM076/prSIM111 | pSIM037 | SacI/NotI |
| pSIM058 | Genomic DNA | prSIM155/prSIM156 | prSIM076/prSIM156 | pSIM037 | SacI/NotI |
| pSIM062 | Genomic DNA | prSIM108/prSIM112 | prSIM076/prSIM112 | pSIM037 | SacI/NotI |
| pSIM081 | pSIM084 | prSIM105/prSIM109 | prSIM076/prSIM109 | pSIM037 | SacI/NotI |
| pSIM091 | Genomic DNA | prSIM105/prSIM109 | prSIM076/prSIM109 | pSIM037 | SacI/NotI |
| pSIM098 | pSIM091 | prSIM197/prSIM109 | prSIM076/prSIM109 | pSIM037 | SacI/NotI |
| pSIM107 | pSIM091 | prSIM202/prSIM109 | prSIM203/prSIM109 | pSIM037 | SacI/NotI |
| pSIM117 | pSIM113 | prSIM202/prSIM109 | prSIM203/prSIM109 | pSIM037 | SacI/NotI |
| pSIM127 | pSIM081 | prSIM202/prSIM109 | prSIM203/prSIM109 | pSIM037 | SacI/NotI |
| pSIM131 | pSIM098 | prSIM202/prSIM109 | prSIM203/prSIM109 | pSIM037 | SacI/NotI |
| pSIM132 | pSIM117 | prSIM202/prSIM109 | prSIM203/prSIM109 | pSIM037 | SacI/NotI |
| pSIM142 | pSIM051 | prSIM249/prSIM110 | N.A. | pFlag-CMV2 | SacI/NotI |
| pSIM143 | pSIM054 | prSIM249/prSIM111 | N.A. | pFlag-CMV2 | SacI/NotI |
| pSIM144 | pSIM058 | prSIM249/prSIM156 | N.A. | pFlag-CMV2 | SacI/NotI |
| pSIM146 | pSIM091 | prSIM249/prSIM109 | N.A. | pFlag-CMV2 | SacI/NotI |
| pSIM147 | pSIM107 | prSIM248/prSIM109 | N.A. | pFlag-CMV2 | SacI/NotI |
| pSIM148 | pSIM107 | prSIM248/prSIM154 | N.A. | pFlag-CMV2 | SacI/NotI |
| pSIM150 | pSIM127 | prSIM248/prSIM109 | N.A. | pFlag-CMV2 | SacI/NotI |
| pSIM154 | pSIM131 | prSIM248/prSIM109 | N.A. | pFlag-CMV2 | SacI/NotI |

| Plasmid | Annealed primers | Backbone | Restriction |
| --- | --- | --- | --- |
| pSIM138 | prSIM239/240 | pX458 | BbsI |
| pSIM139 | prSIM241/242 | pX459 | BbsI |
| pSIM140 | prSIM243/244 | pX458 | BbsI |
| pSIM141 | prSIM245/246 | pX459 | BbsI |

**Supplementary Table S6.**
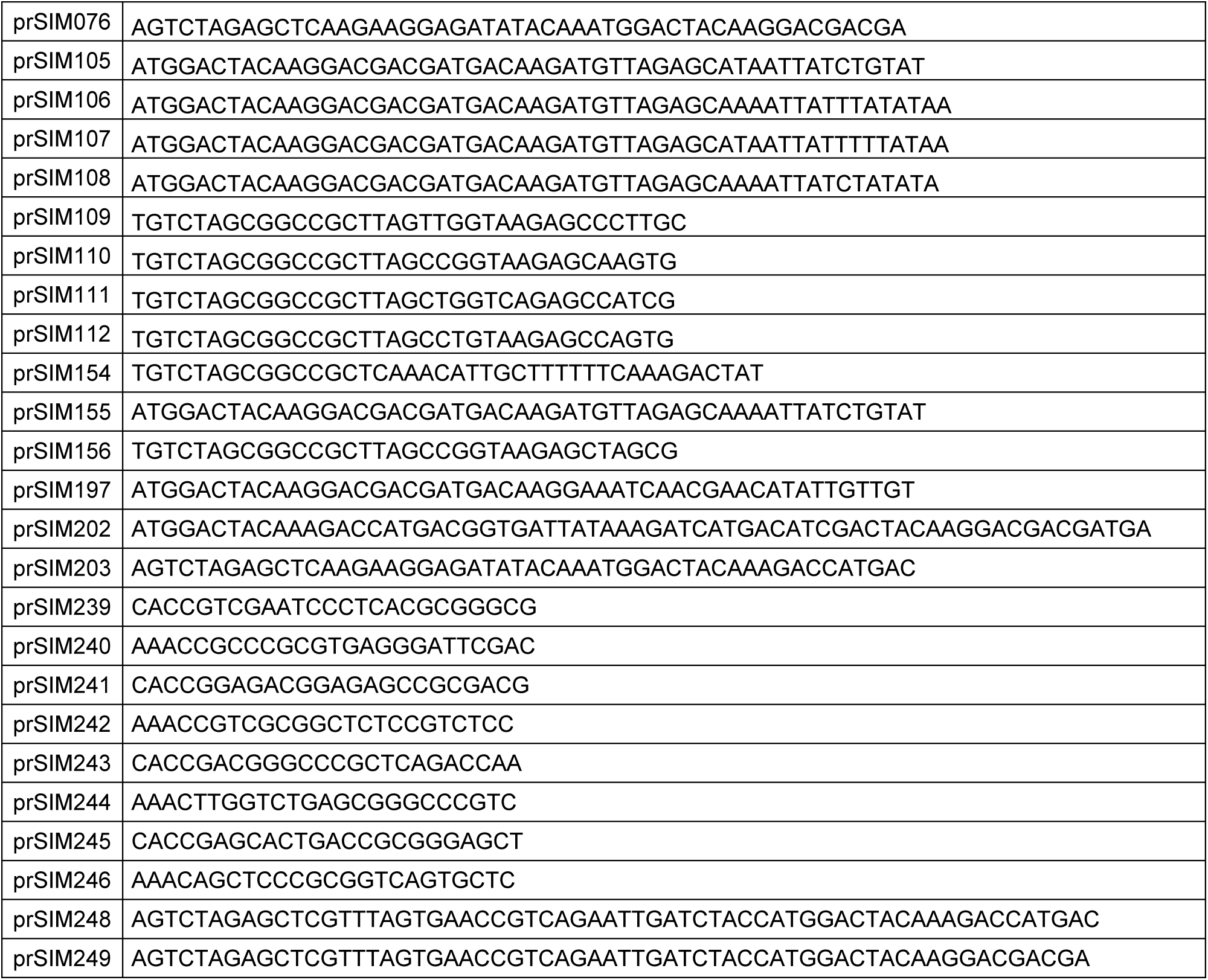
List of primers used in this work

**Supplementary Table S7.** List of antibodies used in this work

| Antibody | Producer (reference) | Final dilution |
| --- | --- | --- |
| Rabbit anti-*Bartonella* | In-house (serum n° 2035) | IF: 1/1000 |
| Rabbit anti-GEF-H1 | ThermoFisher, PA5-32213 | IF: 1/250, WB: 1/500 |
| Rabbit anti-MRCKα | Abcam, ab96659 | WB: 1/500 |
| Rabbit anti-α tubulin | Cell signaling, # 2144 | IF: 1/100 |
| Mouse anti-Flag | Sigma, F1804 | IF/WB: 1/1000 |
| Mouse anti-pMLC | Cell signaling, # 3675 | IF: 1/100 |
| Goat anti-mouse AlexaFluor 488 (secondary) | Jackson Immuno, 115-545-146 | IF: 1/250 |
| Goat anti-mouse AlexaFluor 647 (secondary) | Invitrogen, A21240 | IF: 1/250 |
| Goat anti-rabbit Cy5 (secondary) | Jackson Immuno, 111-175-045 | IF: 1/250 |
| Goat anti-rabbit AlexaFluor 546 (secondary) | Invitrogen, A11035 | IF: 1/250 |
| Donkey anti-rabbit IgG, HRP linked (secondary) | GE Healthcare, NA934 | WB: 1/1000 |
| Sheep anti-mouse IgG, HRP linked (secondary) | GE Healthcare, NA931 | WB: 1/1000 |

